# Selective role of Nck1 in atherogenic inflammation and plaque formation

**DOI:** 10.1101/668129

**Authors:** Mabruka Alfaidi, Christina H. Acosta, Dongdong Wang, James G. Traylor, A. Wayne Orr

## Abstract

While CANTOS established the role of treating inflammation in atherosclerosis, our understanding of endothelial activation at atherosclerosis-prone sites remains limited. Disturbed flow at atheroprone regions primes plaque inflammation by enhancing endothelial NF-κB signaling. Herein, we demonstrate a novel role for the Nck adaptor proteins in disturbed flow-induced endothelial activation. Although highly similar, only Nck1 deletion, but not Nck2 deletion, limits flow-induced NF-κB activation and proinflammatory gene expression. Nck1 knockout mice show reduced endothelial activation and inflammation in both models of disturbed flow and high fat diet-induced atherosclerosis. Bone marrow chimeras confirm that vascular Nck1, but not hematopoietic Nck1, mediates this effect. In contrast, endothelial Nck2 depletion does not affect endothelial activation or atherosclerosis. Domain swap experiments and point mutations identify the Nck1 SH2 domain and the first SH3 domain as critical for flow-induced endothelial activation. We further characterize Nck1’s proinflammatory role by identifying interleukin-1 type I receptor kinase-1 (IRAK-1) as a Nck1-selective binding partner, demonstrating IRAK-1 activation by disturbed flow requires Nck1 *in vitro* and *in vivo*, showing endothelial Nck1 and IRAK-1 staining in early human atherosclerosis, and demonstrating that disturbed flow-induced endothelial activation requires IRAK-1. Taken together, our data reveal a hitherto unknown link between Nck1 and IRAK-1 in atherogenic inflammation.

## Introduction

Atherosclerosis, a chronic lipid-driven arterial inflammatory disease (1), develops at sites of local endothelial activation, a proinflammatory shift in endothelial cell phenotype (2). Local hemodynamic shear stress, a frictional force of blood flow on the endothelium (3), confers protection or susceptibility to endothelial activation, with atheroprotective laminar flow limiting endothelial activation and atheroprone disturbed flow stimulating endothelial activation, characterized by cytoskeletal remodeling, endothelial stiffening, and nuclear factor-kB (NF-κB)-driven proinflammatory gene expression (3-5). Activated endothelial cells recruit monocytes from the circulation, and in the context of hypercholestermia, these monocytes accumulate lipid to drive early fatty streak formation (6). Recruitment of smooth muscle cells from the underlying media contribute to atheroma formation and drive the production of a collagen-rich protective fibrous cap that limits plaque vulnerability to rupture (7). While recent results from the CANTOS trial highlight the important role of limiting inflammation in the treatment of atherosclerosis (8), our understanding of the mechanisms regulating flow-induced endothelial activation remain limited.

The Nck family of adaptor proteins (Nck1 and Nck2) are ubiquitously expressed and share approximately 68% amino acids identity (9).Nck adaptor proteins lack enzymatic activity but control the formation of signaling complexes through three tandem Src homology 3 (SH3) domains and one C-terminal SH2 domain (10). SH2 domains bind with high affinity to tyrosine phosphorylated proteins, whereas the SH3 domains bind to proline-rich sequences (PXXP) in signaling partners, suggesting that Nck serves to couple tyrosine kinase signaling to the activation of downstream pathways (11). The two highly similar Nck proteins are expressed by different genes (11) that play redundant roles during development, as deletion of both Nck isoforms results in an embryonic lethal phenotype due to impaired vasculogenesis while deletion of only one isoform did not (9). While Nck1 and Nck2 play redundant roles in regulating angiogenesis in mouse models of retinopathy (12), Nck2 may play a dominant role in PDGF-induced actin polymerization in NIH3T3 cells (13). In contrast, Nck1 plays a more dominant role in T cell receptor-induced ERK activation (14), suggesting non-compensating roles during phenotypic regulation post-development. However, the signaling effects of Nck1/2 outside the context of cytoskeleton remodeling are incompletely understood.

We have previously demonstrated that treatment with a membrane-permeable peptide corresponding to a Nck-binding PXXP sequence blunts flow-induced endothelial activation *in vitro* and significantly reduces both inflammation and vascular permeability at atheroprone areas exposed to disturbed flow *in vivo* (15). However, this peptide could be reducing atherosclerotic inflammation through off-target effects on other SH3 containing adaptor proteins. Furthermore, the anti-inflammatory effects observed *in vivo* could be due to effects of this peptide on other cell types besides endothelial cells (16). Interestingly, the Nck1 gene locus at chromosome 3q22.3 has been associated with atherosclerosis susceptibility and myocardial infarction (17), and a genome wide association study identified Nck1 as a novel coronary artery disease susceptibility loci (18). Herein we utilized both cell culture and animal models of disturbed blood flow and high fat diet induced atherosclerosis to characterize the non-compensating functions of Nck1 and Nck2 in disturbed flow-induced endothelial activation.

## Results

### Nck1/2 deletion and their effects on shear stress-induced endothelial activation

To investigate the direct roles of Nck1/2 in shear stress induced endothelial activation, we adopted a loss of function model where the cellular levels of Nck1 and Nck2 were reduced by siRNA transfection. The efficiency of transfection was confirmed using Western blotting, with a 70% knockdown of Nck1 and 84% knockdown of Nck2 (Figure 1A). Since the acute endothelial response to shear stress models the sustained signaling observed with chronic disturbed flow (19), endothelial cells were subjected to acute shear stress for 0, 5, 15, or 30 minutes, and the early signaling effects were measured. NF-κB activation was assessed by measuring p65 S536 phosphorylation or nuclear translocation. Nck1/2 knockdown cells showed a significant reduction in both NF-κB phosphorylation (Figure 1B-C) and NF-κB nuclear translocation (55± 7.8% mock vs. 13±1.4% in Nck1/2 siRNA, p<0.0001) (Figure 1D-E). To confirm the combined effect of Nck1/2 knockdown, we used CRISPR/Cas9 editing to generate a stable human aortic endothelial cell (HAEC) line lacking in both Nck1 and Nck2. Nck1/2 double knock out cells (Nck1/2 DKO) show a similar decrease in shear stress-induced NF-κB phosphorylation (Figure 1F-G) and nuclear translocation (Figure1H-I) compared to scrambled controls (Figure 1F-I). Similar results were also observed in mouse aortic endothelial cells (MAECs) isolated from endothelial-specific Nck1/2 double knockout mice (iEC-Nck1/2 DKO; VE-cadherinCreERT2^tg/?^, Nck1^−/−^, Nck2^fl/fl^, ApoE^−/−^) (Supplemental Figure 1A-C).

**Figure 1.**
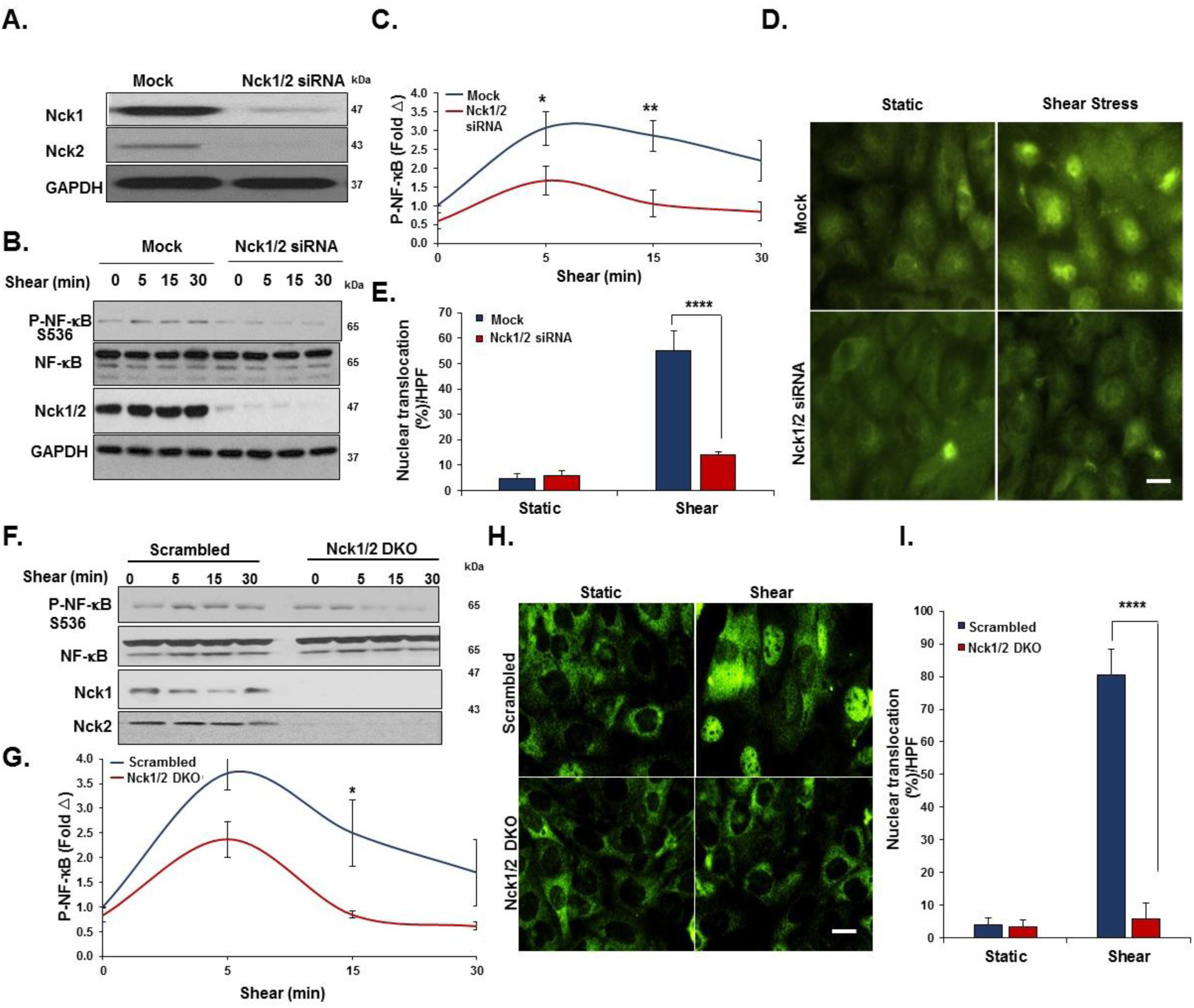
Nck1/2 deletion ameliorate acute shear stress induced NF-κB activation. **A)** Human aortic endothelial cells (HAECs) were transfected with siRNA specific for Nck1/2, and transfection efficiency was assessed using Western blot. **B-C)** HAECs were subjected to acute shear stress for the indicated times, and NF-κB activation was assessed by detection of p65 serine 536 phosphorylation using Western blotting. **D-E)** p65 nuclear translocation was measured after 45 minutes of shear stress in Nck1/2 siRNA and mock control cells. **F-I)** Nck1/2 was deleted from HAECs using CRISPR/Cas9 editing, and shear stress-induced NF-κB activation was assessed by **(F-G)** Western blotting for p65 phosphorylation and **(H-I)** staining for p65 nuclear translocation. Densitometric analysis was performed using Image j. Images were analyzed using NIS Elements software. Scale bars=50µm. Data are mean ± SEM, n=4, analyzed by 2-Way ANOVA followed by Bonferroni’s post-test, *p<0.05, **p<0.01, ****p<0.0001.

In response to chronic oscillatory shear stress (OSS), NF-κB activation drives proinflammatory gene expression, including ICAM-1 and VCAM-1 (20). We found that siRNA-mediated Nck1/2 depletion reduced oscillatory shear stress-induced VCAM-1 and ICAM-1 protein expression (Figure 2A/B). Compared to the mock controls, Nck1/2 siRNA depleted cells showed significantly less NF-κB activation (Figure 2B) and mRNA expression of ICAM-1 and VCAM-1 (Figure 2C). Shear stress-induced expression of the atheroprotective gene KLF2 was not affected by Nck1/2 depletion (Figure 2C). Consistent with these data, Nck1/2 DKO endothelial cells show a similar reduction in oscillatory shear stress-induced NF-κB activation (Figure 2D/E) and VCAM-1/ICAM-1 expression (Figure 2D-F). Taken together, these data suggest that Nck1/2 expression is required to couple atheroprone hemodynamics to NF-κB activation and proinflammatory gene expression.

**Figure 2.**
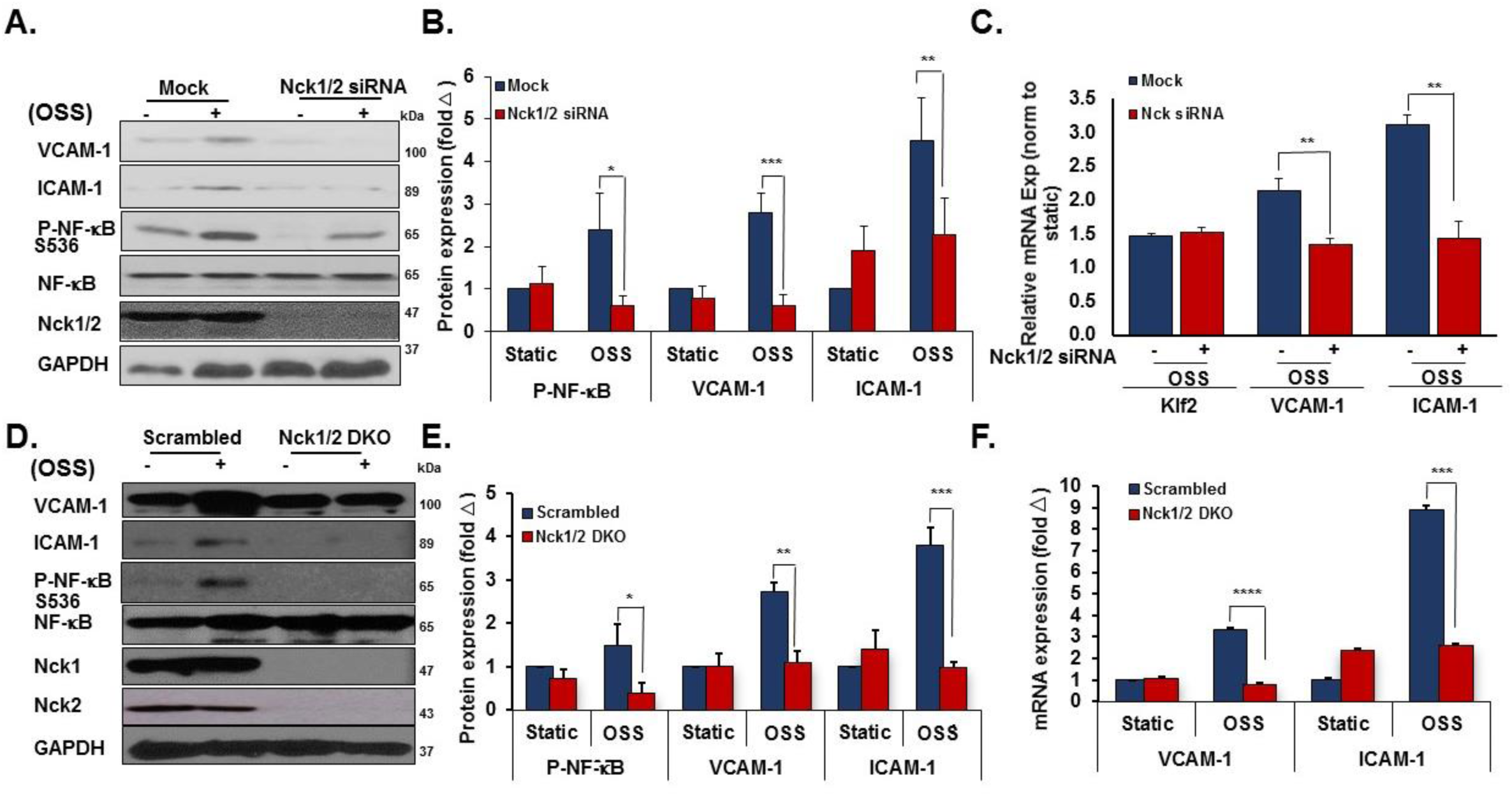
Nck1/2 deletion ameliorate chronic oscillatory shear stress induced endothelial activation. **A/B)** HAECs were transfected with Nck1/2 siRNA, and oscillatory shear stress (OSS; ±5 dynes/cm^2^ with 1 dyne/cm^2^ forward flow)-induced proinflammatory gene expression (VCAM-1, ICAM-1) and proinflammatory signaling (P-NF-κB Ser536) was assessed by Western blotting. **C)** HAECs were treated as in **(A)**, and mRNA expression was assessed by qRT-PCR. **D/E)** Nck1/2 was deleted from HAECs using CRISPR/Cas9 and OSS-induced proinflammatory gene expression and signaling was assessed by Western blotting. **F)** HAECs were treated as in **(D)** & **(A)**, and mRNA expression was assessed by qRT-PCR. Data are mean ± SEM, n=4, analyzed by 2-Way ANOVA followed by Bonferroni’s post-test, *p<0.05, **p<0.01, ***p<0.001, ****p<0.0001.

### Deletion of Nck1, but Not Nck2, ameliorates shear stress-induced endothelial activation

Even though they are expressed by different genes, Nck1 and Nck2 proteins share a high sequence identity (68% overall) (10) and their functions are generally regarded as overlapping (21). However, emerging evidence has suggested the independent contribution of the two isoforms in a variety of responses, including T cell activation, cytokinesis, and podocyte cytoskeletal dynamics (22-24). To investigate the selective roles of Nck1 and Nck2, we utilized Nck1 and Nck2 selective siRNAs that result in a 75% and 85% knockdown, respectively without affecting the expression of the other isoform (Figure 3A). In response to shear stress, Nck1 depleted cells showed significantly less NF-κB p65 phosphorylation (Figure 3B/C) and nuclear translocation (Figure 3D/E), whereas Nck2 depletion did not affect NF-κB activation by flow. To confirm these effects, we utilized lentiviral shRNA constructs to selectively deplete Nck1 (100% knockdown) and Nck2 (90% knockdown) (Figure 3F). Similar to siRNA data, only HAECs expressing Nck1 shRNA showed significant amelioration of NF-κB phosphorylation (Figure 3G/H) and nuclear translocation (Figure 3I/J), whereas cells expressing Nck2 shRNA did not significantly differ from cells expressing scrambled shRNA. MAECs isolated from Nck1 KO mice showed similar results with remarkable reduction in NF-κB activation following shear stress, whereas MAECs from iEC-Nck2 KO mice showed the usual shear stress-induced NF-κB activation (Supplemental Figure 2A-C). To assess the specific role of Nck1 in atheroprone disturbed flow models, HAECs transfected with Nck1 and Nck2 siRNA or shRNA were exposed to oscillatory shear stress for 18 hours, and proinflammatory signaling (NF-κB) and proinflammatory gene expression (VCAM-1/ICAM-1) were assessed. Oscillatory flow-induced NF-κB activation (p65 Ser536 phosphorylation) and VCAM-1/ICAM-1 protein expression (Figure 4A/B) and mRNA levels (Supplemental Figure 3A) were blunted by Nck1 siRNA and Nck1 shRNA (Figure 4D/E), (Supplemental Figure 3B), whereas Nck2 depletion (siRNA and shRNA) had no significant effects on any of these responses.

**Figure 3.**
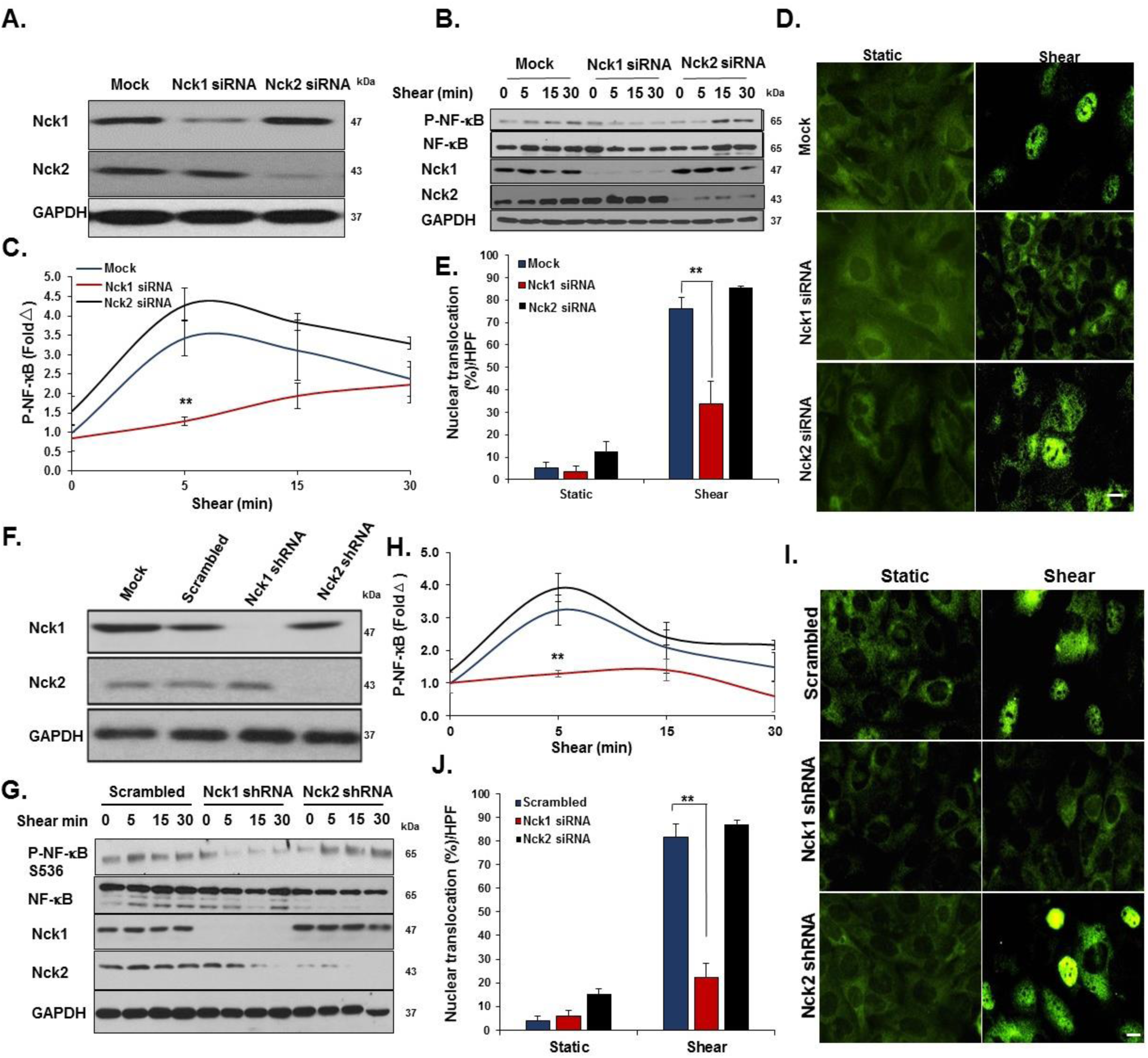
Nck1, but not Nck2, deletion ameliorates acute shear stress induced activation. **A)** Transfection efficiency of selective Nck1 and Nck2 knockdown in HAECs using siRNA. **B-E)** HAECs lacking either Nck1 or Nck2 were subjected to acute shear stress for the indicated times, and NF-κB activation was assessed by measuring **(B/C)** NF-κB phosphorylation and **(D/E)** nuclear translocation. **F)** Lentiviral Nck1 and Nck2 knocking down by shRNA. **G-H)** HAECs expressing either Nck1 shRNA or Nck2 shRNA were subjected to acute shear stress for the indicated times, and NF-κB activation was assessed by measuring **(G/H)** NF-κB phosphorylation and **(I/J)** nuclear translocation. Scale bars=50µm. Data are mean ± SEM, n=4, analyzed by 2-Way ANOVA followed by Bonferroni’s post-test, **p<0.01.

**Figure 4.**
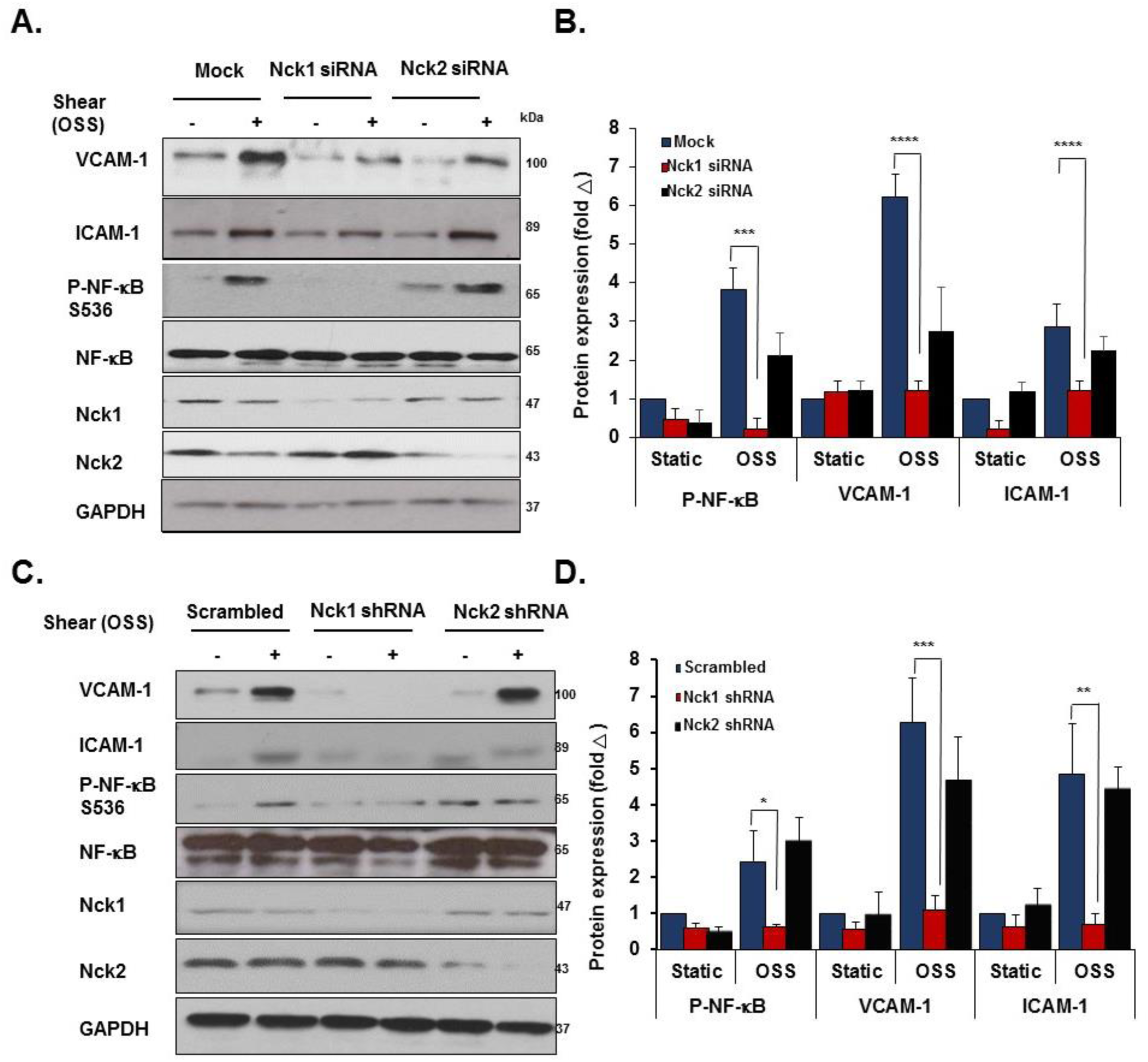
Nck1, but not Nck2, deletion ameliorates chronic oscillatory shear stress induced activation. **A/B)** HAECs were transfected with either Nck1 or Nck2 siRNA, and oscillatory shear stress (OSS, 18h) induced proinflammatory gene expression (ICAM-1/VCAM-1) and signaling (P-NF-κB Ser536) were assessed by Western blotting. **C/D)** HAECs were transfected with either Nck1 or Nck2 shRNA, and oscillatory shear stress (OSS, 18h) induced proinflammatory gene expression (ICAM-1/VCAM-1) and signaling (P-NF-κB Ser536) were assessed by Western blotting. Data are from n=4, mean ± SEM, analyzed by 2-Way ANOVA followed by Bonferroni’s post-test, *p<0.05, **p<0.01, ***p<0.001, ****p<0.0001.

While these data identify Nck1 as a critical regulator of NF-κB activation and proinflammatory gene expression by atheroprone hemodynamics, Nck1 did not affect all shear stress responses. Neither Nck1 nor Nck2 depletion affected KLF2 expression under oscillatory shear stress (Supplemental Figure 3) or activation of other classic shear stress-induced signaling pathways, such as Akt, eNOS, and ERK1/2 phosphorylation (Supplemental Figure 4). Collectively these data demonstrate a critical role for Nck1 in endothelial activation by atheroprone flow, whereas Nck2 has no effect.

### Nck1 mediates disturbed flow-induced endothelial activation in vivo

Having shown that Nck1 regulates endothelial activation by atheroprone flow *in vitro*, we sought to investigate the differential effects of Nck1 and Nck2 in an *in vivo* model of disturbed flow. Following tamoxifen induction, inducible endothelial-specific control mice (iEC-Control; VE-cadherinCreERT2^tg/?^, ApoE^−/−^), Nck1 knockout (VE-cadherinCreERT2^tg/?^, Nck1^−/−^, ApoE^−/−^), endothelial-specific Nck2 knockouts (iEC-Nck2 KO; VE-cadherinCreERT2^tg/?^, Nck2^fl/fl^, ApoE^−/−^), and endothelial-specific Nck1/2 double knockout mice (iEC-Nck1/2 DKO; VE-cadherinCreERT2^tg/?^, Nck1^−/−^, Nck2^fl/fl^, ApoE^−/−^) were subjected to partial carotid ligation (PCL) to induce disturbed flow-associated endothelial activation specifically in the left carotid artery (25, 26). Changes in endothelial mRNA expression was assessed after 48 hours, whereas changes in inflammatory gene expression and macrophage recruitment was assessed after 7 days (Figure 5A). To assess endothelial activation, endothelial mRNA was isolated from the left and right carotid vessels by TRIzol flush after tissue harvesting (27). The purity of intimal mRNA and medial/adventitial mRNA was confirmed by measuring platelet endothelial cell adhesion molecule-1 (PECAM-1) and α-smooth muscle actin (SMA) expression (Supplemental Figure 5A/B). Furthermore, endothelial-specific deletion of Nck2 was confirmed in iEC-Nck2 KO and iEC-DKO mice, as Nck2 mRNA expression was depleted in the intimal but not the medial/adventitial fractions (Supplemental Figure 5C/D). KLF2 showed decreased expression in the ligated left carotid compared to the right carotid control, confirming a OSS flow-associated gene expression profile (Figure 5B). However, this downregulation did not differ among experimental animals. Nck1 knockouts showed a pronounced reduction in oscillatory flow-induced VCAM-1 and ICAM-1 mRNA expression (Figure 5B). However, Nck2 deletion (iEC-Nck2 KO) did not affect disturbed flow-induced VCAM-1 and ICAM-1 expression, and VCAM-1/ICAM-1 mRNA expression did not significantly decrease in iEC-Nck1/2 DKO mice compared to Nck1 KO mice (Figure 5B).

**Figure 5.**
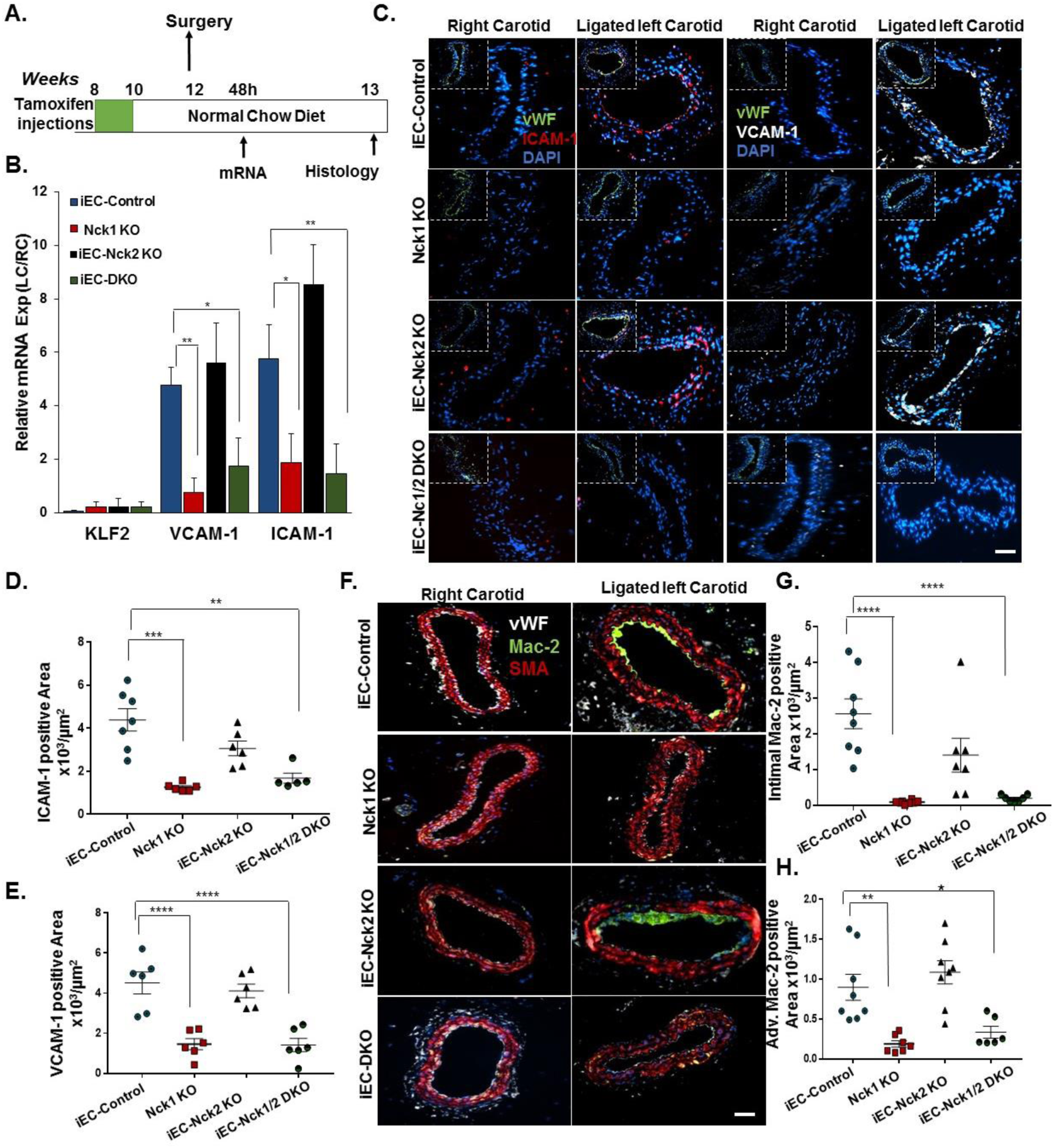
Ablation of Nck1 but not Nck2 blunts Partial Carotid Ligation Induced Inflammation. **A)** Schematic of the study in which four groups of mice were subjected to the ligation surgery as indicated animal genotypes and time of surgery. **B)** Endothelial mRNA analysis from iEC-Control, Nck1 KO, iEC-Nck2 KO, and iEC-Nck1/2 DKO mice. mRNA from the ligated left carotid was normalized to the unligated right carotid and to the housekeeping gene ß-microglobulin. Data analyzed by 2-Way ANOVA and Bonferroni’s post-test, *p<0.05, **p<0.01. **C-E)** ICAM-1 (red) and VCAM-1 (white) in the ligated left carotid compared to the unligated right carotid arteries among experimental groups. Endothelial cells were stained with Von Willebrand factor (vWF) and the nuclei counterstained with DAPI. **F-H)** Macrophage staining (Mac-2, green) in the ligated and the unligated carotid arteries among experimental groups. Smooth muscle cells (α-smooth muscle actin (SMA), red), and endothelium (vWF, white). Scale bars=100µm. Images analyzed using Nis Elements software, from n=7-10 mice/group. Data are mean ± SEM, analyzed by 1-Way ANOVA and Tukey’s post-test, *p<0.05, **p<0.01, ***p<0.001, ****p<0.0001.

To examine early atherogenic remodeling in the ligated carotid arteries, tissues were collected 7 days post-ligation and assessed for markers of inflammation by immunohistochemistry. Consistent with early changes in mRNA expression, VCAM-1 and ICAM-1 protein levels were significantly reduced in Nck1 KO mice after PCL compared to iEC-Control mice (Figure 5 C-E) whereas iEC-Nck2 KO animals were similar to controls. Similarly, Nck1 KO mice showed a significant reduction in intimal macrophage recruitment (Mac2-positive area) compared to iEC-Control mice (Figure 5F/G). Nck1 KO also significantly reduced adventitial macrophage content (Figure 5F/H), but endothelial Nck2 deletion (iEC-Control vs iEC-Nck2 KO; Nck1 KO vs iEC-Nck1/2 DKO) did not affect intimal or adventitial macrophage infiltration (Figure 5F-H). Taken together, our data suggest a direct role for Nck1 in regulating endothelial activation and macrophage recruitment under atheroprone hemodynamics.

### Nck1 deletion reduces atherosclerotic plaque formation

Atheroprone flow establishes local susceptibility to endothelial activation and to diet-induced atherosclerotic plaque development (28). Having observed differential effects for Nck1 and Nck2 in response to atheroprone flow-induced endothelial activation, we sought to determine if atherosclerotic plaque development was altered in Nck1 KO mice. iEC-Control, Nck1 KO, iEC-Nck2 KO, or iEC-DKO mice were fed high fat diet (HFD) for 12 weeks to induce spontaneous atherosclerosis. No significant differences were observed in body weight over the 12 weeks of HFD feeding, though Nck1 KO mice tended to be smaller (Supplemental Figure 6A), and no changes were noted for heart weight (Supplemental Figure 6B) or for plasma cholesterol, triglycerides or HDL levels among the experimental groups (Supplemental Figure 6C-E). However, Nck1 KO mice showed significant reductions in the plasma levels of several proinflammatory mediators, including interleukin-1α (IL-1α), IL-1ß, TNF-α, and MCP-1 (Supplemental Figure 7), highlighting the proinflammatory role of Nck1. Atherosclerotic lesion formation was assessed in four different vascular sites, including the aorta, the aortic roots, the brachiocephalic artery, and the right and left carotid sinuses. *En face* analysis of atherosclerosis in the aorta was assessed by Oil red O staining and calculated as the percent lesion area compared to the total surface area of the aorta (Figure 6A). While iEC-Nck2 KO mice did not differ from iEC-Controls, Nck1 KO mice show a significant reduction in plaque burden in the aorta (Figure 6A/B), the carotid sinus (Figure 6C/D) and brachiocephalic artery (Figure 6H/I). Atherosclerosis did not decrease further in the iEC-Nck1/2 DKO compared to the Nck1 KO, suggesting that Nck1 is the key regulator and the endothelial Nck2 does not significantly contribute to atherogenic endothelial activation.

**Figure 6.**
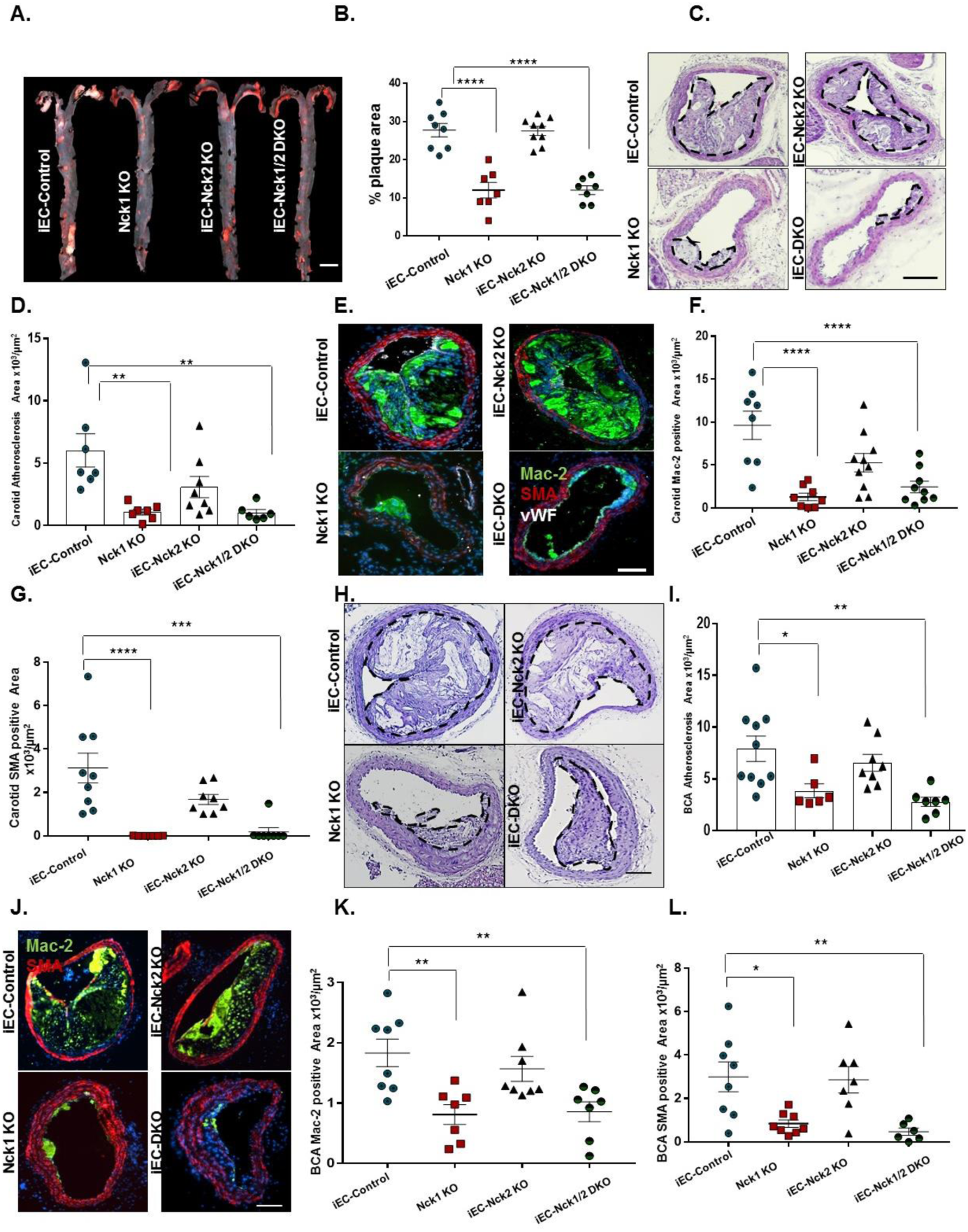
Global deletion of Nck1 but not Nck2 blunts high fat diet-induced atherosclerosis. iEC-Control, Nck1 KO, iEC-Nck2 KO, and iEC-Nck1/2 DKO mice were fed high fat diet (HFD) for 12 weeks. **A)** Representative *en face* morphometric images of total aortic lesion area and **(B)** calculated whole aortic atherosclerosis (% of the total surface area). Scale bar=1mm. **C)** Representative stained Hematoxylin and Eosin (H&E) images of carotid atherosclerosis and **(D)** quantification of carotid atherosclerotic area among experimental groups. **E-G)** Analysis of carotid plaque cellular content following staining for macrophages (Mac-2, green), smooth muscle cells (α-smooth muscle actin (SMA), red), and endothelium (vWF, white). **H)** H&E staining of brachiocephalic arteries (BCA) after HFD feeding in iEC-Controls, Nck1 KO, iEC-Nck2 KO, iEC-DKO mice. **I)** Quantification of atherosclerosis burden in brachiocephalic arteries among experimental groups. **J)** Plaque composition assessed by staining for macrophages (Mac2 positive) and smooth muscle cells (SMA positive) in brachiocephalic lesions. **K)** Quantification of macrophage (Mac2 positive) and **(L)** SMA positive areas in brachiocephalic arteries. Data are mean ± SEM, n=6-10/group. Data analyzed by 1-Way ANOVA and Tukey’s post-test, *p<0.05, **p<0.01, ***p<0.001, ****p<0.0001. Scale bars=100µm.

To assess atherosclerotic plaque characteristics in this model, plaques were stained for macrophage (Mac2 positive area) and smooth muscle (a-smooth muscle actin (SMA) positive) levels. Compared to iEC-Control mice, Nck1 KO mice show a significant reduction in macrophage area in both the carotid sinus (Figure 6E/F) and brachiocephalic arteries (Figure 6J/K). Similarly, Nck1 deletion reduces plaque smooth muscle (SMA positive) area (Figure 6G/L) and lipid core area (Supplemental Figure 8A) at these sites, consistent with the very early stages of plaque formation observed in Nck1 KO and iEC-Nck1/2 DKO mice. Atherosclerotic lesions in the brachiocephalic artery and carotid sinuses tend to be less well developed than the plaques in the aortic root, potentially due to delayed onset of plaque formation at these sites (29). Unlike other sites, we did not observe any differences in plaque size in the aortic root (Supplemental Figure 8B). However, the plaques that formed showed reduced macrophage area (Supplemental Figure 8C/D) and enhanced smooth muscle area (Supplemental Figure 8C/E), suggestive of enhanced plaque stability.

### Deletion of Nck1 confers atheroprotection in resident vascular bed cells compared to myeloid linage cells

To determine the relative contribution of Nck1 deletion in either vascular cells or myeloid cells to atherosclerosis formation, we performed bone marrow transplantation experiments where Nck1 KO or Nck1 WT bone marrow was transplanted into irradiated Nck1 KO or Nck1 WT recipients. Successful chimera were confirmed with the alteration in Nck1 expression in the myeloid lineage cells (Supplemental Figure 9). After 12 weeks of HFD, there were no significant changes in neither body weights (Supplemental Figure 10A) nor plasma cholesterol levels (Supplemental Figure 10B) in mice lacking Nck1 expression in myeloid cells compared to other experimental groups. Consistent with our *in vitro* observations on endothelial activation, Nck1 KO mice still showed significant reduction in aortic atherosclerosis when reconstituted with Nck1 WT bone marrow, whereas Nck1 WT mice receiving bone marrow from Nck1 KO mice did not show any significant difference compared to Nck1 WT mice receiving Nck1 WT bone marrow (Figure 7A/B). Similarly, carotid (Figure 7C/D) and brachiocephalic atherosclerotic (Figure 7H/D) lesions were two to three-fold less in Nck1 KO mice receiving Nck1 WT bone marrow compared to controls (p<0.0001, p<0.05, respectively). Cross-sectional analysis of mac-2 and SMA positive areas within carotids (Figure 7E-G) and BCA (Figure 7J-L) lesions showed significant reductions of carotid mac-2 and SMA areas only when Nck1 was depleted in the resident vascular cells but not when Nck1 was depleted in the bone marrow. These data support the role of vascular wall Nck1 in atherogenic inflammation and atherosclerotic plaque development and suggest that myeloid Nck1 expression does not contribute to atherosclerotic plaque formation.

**Figure 7.**
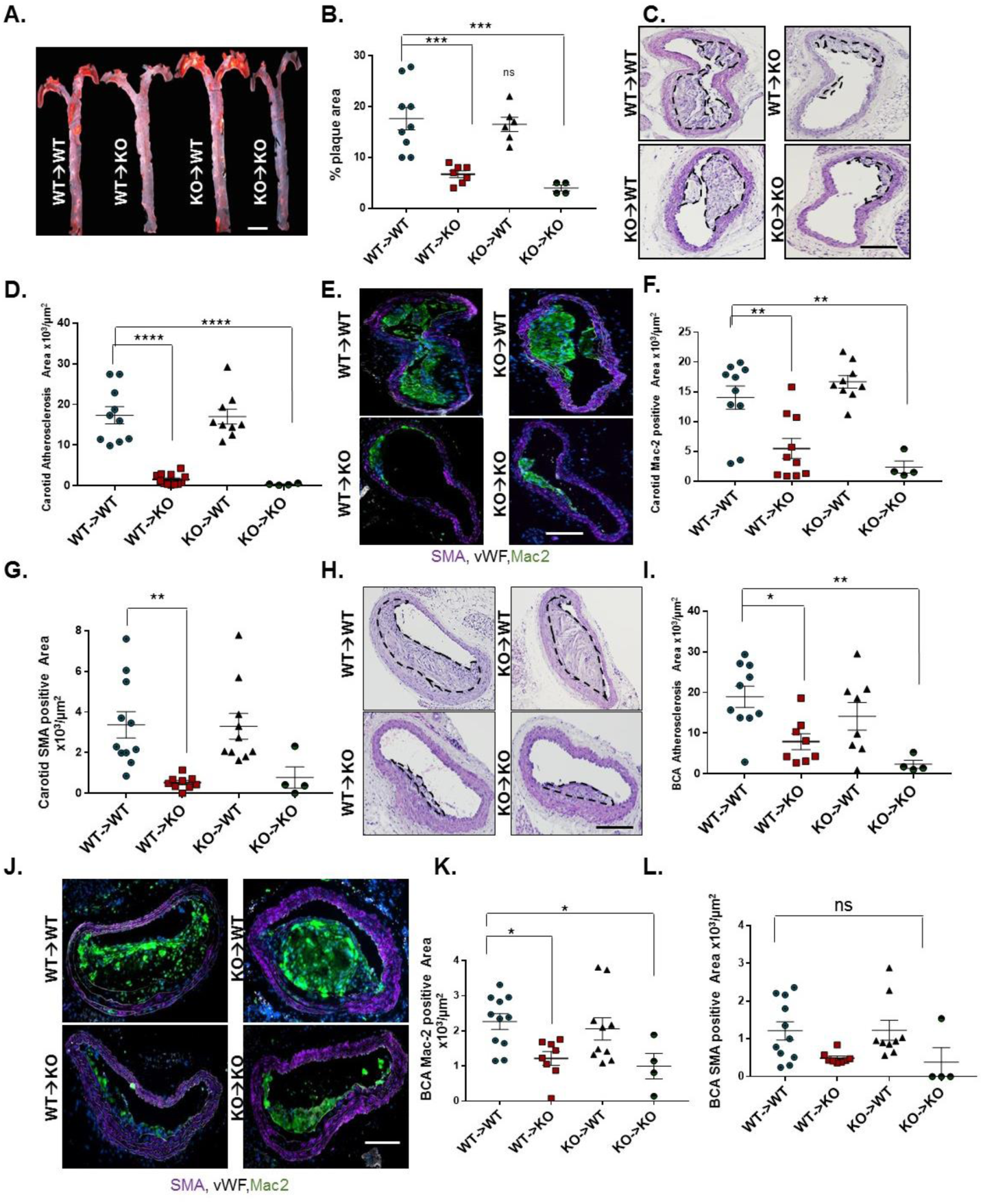
Nck1 deletion from vascular wall but not hematopoietic cells has atheroprotection effect. Nck1 wild-type (WT) and Nck1 KO mice were irradiated and received bone marrows from either Nck1 WT or Nck1 KO. Two weeks after irradiation, the four groups of mice were fed HFD for 12 weeks. **A)** Representative *en face* morphometric images of total aortic lesion area and **(B)** calculated whole aortic atherosclerosis (% of the total surface area). Scale bar=1mm. **C)** Representative stained H&E images of carotid atherosclerosis and **(D)** quantification of carotid atherosclerotic area among experimental groups. **E-G)** Analysis of plaque cellular content following staining for macrophages (Mac-2, green), smooth muscle cells (α-smooth muscle actin (SMA), purple), and endothelium (vWF, white). **H)** H&E staining of brachiocephalic arteries (BCA) after HFD feeding in WT→WT, WT→KO, KO→WT, Nck1 KO→Nck1 KO mice. **I)** Quantification of atherosclerosis burden in brachiocephalic arteries among experimental groups. **J)** Plaque composition assessed by staining for macrophages (Mac2 positive) and smooth muscle cells (SMA positive) in brachiocephalic lesions. **K)** Quantification of macrophage (Mac2 positive) and **(L)** SMA positive areas in brachiocephalic arteries. Data are mean ± SEM, n=4-11/group. Data analyzed by 1-Way ANOVA and Tukey’s post-test, *p<0.05, **p<0.01, ***p<0.001, ****p<0.0001. ns, non-significant. Scale bars=100µm.

### Nck1 exerts the non-redundant effects on endothelial activation via its SH2 domain

To mechanistically understand why Nck1 but not the highly homologous Nck2 regulates endothelial activation, we conducted domain swap experiments mixing the Nck1 SH2 domain with Nck2 SH3 domains and the Nck2 SH2 domain with Nck1 SH3 domains (Figure 8A). We confirmed similar expression levels of transfected constructs encoding Nck1, Nck2, the Nck1 SH2/Nck2 SH3 chimera, and the Nck2 SH2/Nck1 SH3 chimera in Nck1/2 DKO cells by Western blotting (Figure 8B). Nck1, but not Nck2, re-expression restored oscillatory shear stress-induced NF-κB activation (Figure 8C), VCAM-1 expression (Figure 8D) and ICAM-1 expression (Supplemental Figure 11) in the Nck1/2 DKO cells. However, only the Nck1 SH2/Nck2 SH3 chimera showed a similar restoration, suggesting that the Nck1 SH2 domain is essential to form the signaling complex required for oscillatory flow-induced endothelial activation. However, the redundancy of the Nck1 and Nck2 SH3 domains does not exclude them as important for the activation of this response. To gain further insight into the Nck1 domains regulating oscillatory flow-induced NF-κB activation, we introduced single point mutations (Figure 8E/F) to inactivate the Nck1 SH2 domain (R308M, Nck1 SH2*), the first SH3 domain (W38K, Nck1 SH3.1*), the second SH3 domain (W143K, Nck1 SH3.2*), or the third SH3 domain (W229K, Nck1 SH3.3*) (30). Following oscillatory shear stress exposure, NF-κB activation and VCAM-1 expression were only blunted in cells expressing the Nck1 SH2* and the Nck1 SH3.1* constructs (Figure 8G-I), suggesting critical roles for Nck1 SH2-based phosphotyrosine binding and Nck1 SH3.1-based binding partners.

**Figure 8.**
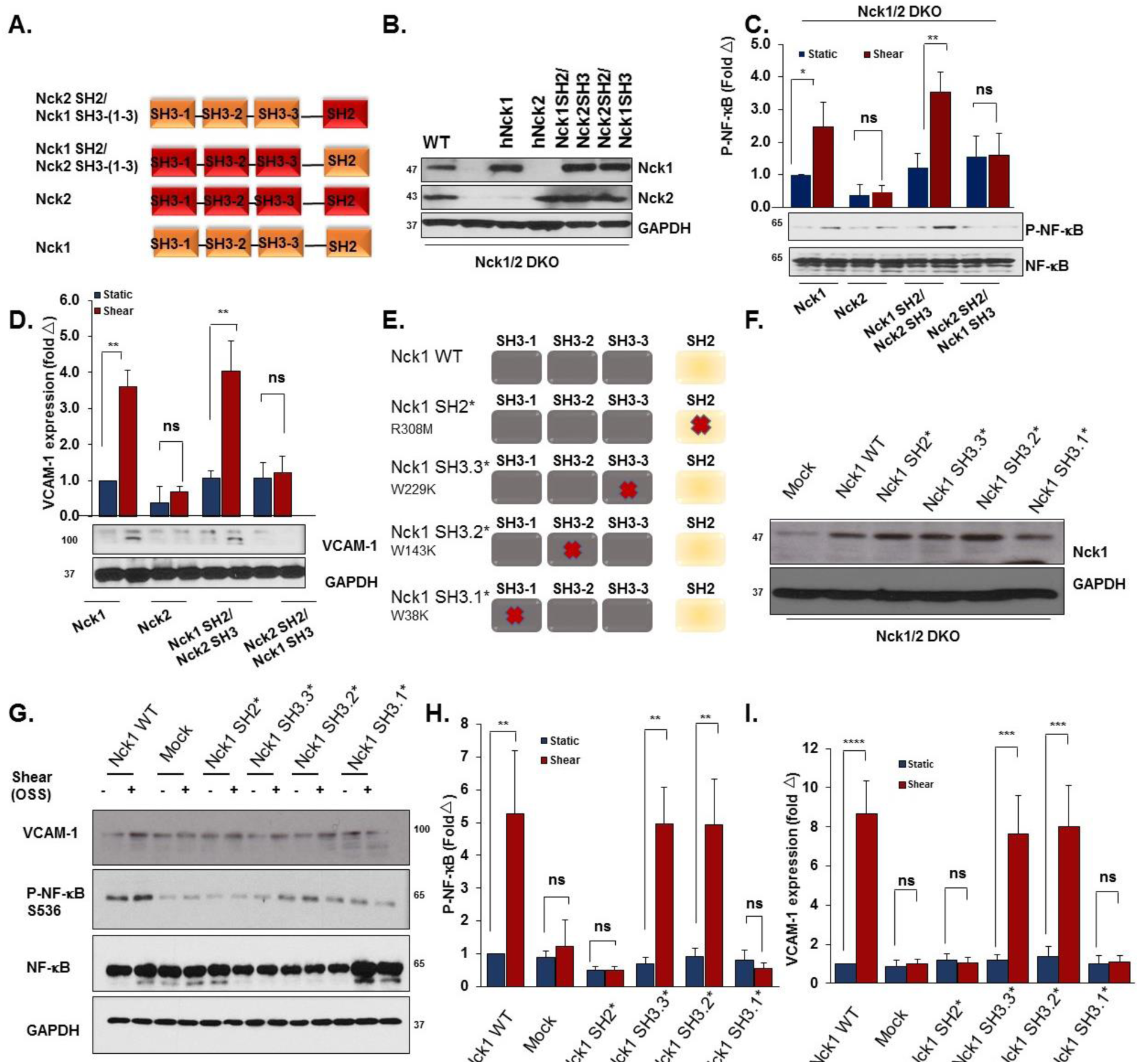
Nck1 Regulates Shear Stress Induced Inflammation via its SH2 domain and the first domain of its SH3 domains. **A)** Schematic showing the domain structure of Nck1 and Nck2 and the two chimeras of Nck1 SH2/ Nck2 SH3 (1-3) and Nck2 SH2/ Nck1 SH3 (1-3). **B)** Western blot analysis showing comparable transduction efficiency of Nck1/2 chimeras following introducing the constructs in Nck1/2 DKO cells. n=3. **C-D)** Nck1/2 DKO HAECs were transduced with constructs in **(A)**, and oscillatory shear stress-induced proinflammatory signaling (P-NF-κB Ser536) and gene expression (VCAM-1) were assessed. **E)** Schematic of Nck1 domain point mutations, and **(F)** Western blot analysis showing the efficiency of re-expression of different Nck1 mutants in Nck1/2 DKO HAECs. **G-I)** Nck1/2 DKO HAECs were transiently transfected with Nck1 or Nck1 variants described in **(E)**, and OSS-induced proinflammatory signaling and gene expression assessed as indicated above. Data are from n=4, represented as mean ± SEM, analyzed by 2-Way ANOVA, and Bonferroni’s post-test, *p<0.05, **p<0.01, ***p<0.001, ****p<0.0001.

### Nck1 interacts with interleukin-1 type I receptor kinase-1 (IRAK-1) in response to atheroprone hemodynamics

While these data suggest a critical role for Nck1 in atherogenic endothelial activation, the Nck1 binding partner involved remains unknown. A proteomics study by Jacquet *et al* identified IRAK-1 as a Nck1-specific SH2 interacting partner (24). IRAK-1, a serine/threonine kinase extensively investigated in the interleukin-1 signaling pathway, is required for interleukin-1-induced NF-κB activation (31). Therefore, we exposed human endothelial cells to either atheroprotective laminar or atheroprone oscillatory shear stress, and assessed activation of IRAK-1 (Ser/Thr209 phosphorylation). While we did not observe any significant changes in Nck1 and IRAK-1 total protein levels, cells exposed to atheroprone OSS showed marked induction of IRAK-1 phosphorylation (Supplemental Figure 12), suggesting that OSS promotes IRAK-1 activation. To test the hypothesis that IRAK-1 phosphorylation requires Nck1, IRAK-1/Nck1 interactions were assessed by co-immunoprecipitation in endothelial cells exposed to flow. Following shear stress exposure, Nck1 interactions with IRAK-1 were significantly enhanced (Figure 9A-C), whereas Nck2 did not interact with IRAK-1 (Figure 9A), confirming the differences between Nck1 and Nck2 in shear stress signaling. Moreover, Nck1-depleted HAECs subjected to OSS showed a marked reduction in IRAK-1 phosphorylation (Supplemental Figure 13). Following partial carotid ligation *in vivo*, phospho-IRAK-1 staining is markedly enhanced in the ligated carotids, predominantly within the endothelium (Figure 9D). However, endothelial cell IRAK-1 phosphorylation was significantly abolished in Nck1 KO and Nck1/2 DKO mice (Figure 9D/E). Consistent with this data, BCA atherosclerotic plaques showed enhanced phospho-IRAK-1 staining in the endothelium that was markedly reduced in Nck1 KO mice (Supplemental Figure 14), confirming the role of Nck1 in IRAK-1 activation at atherosclerosis prone sites.

**Figure 9.**
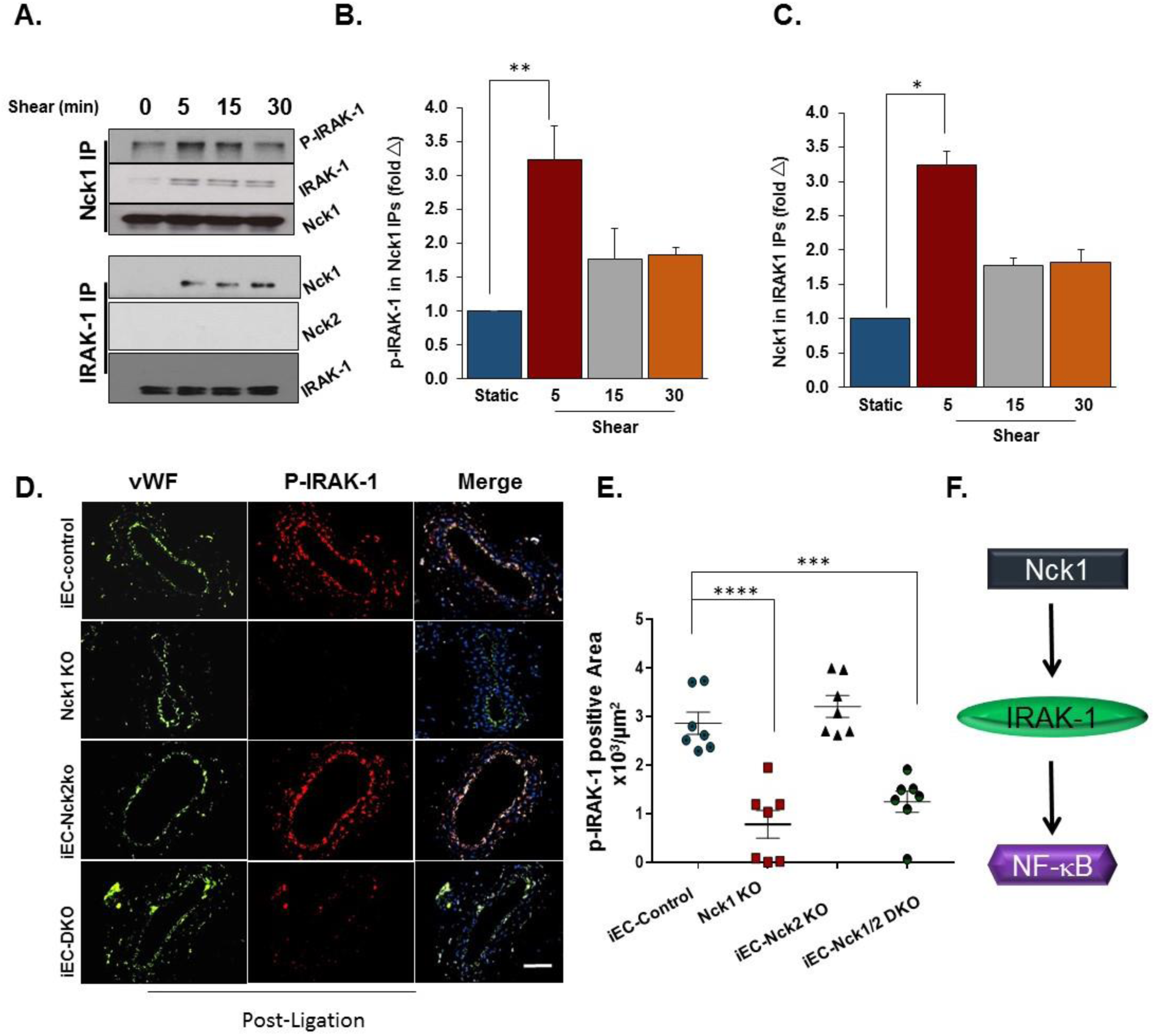
Nck1 interacts with IRAK1and IRAK1 activation requires Nck1 in response to shear stress. **A)** Immunoblotting showing Nck1 and IRAK-1 co-immunoprecipitations (Co-ip) and confirms Nck1 and IRAK-1 interactions and that is shear stress-dependent. **B-C)** Graphical representations to the levels of p-IRAK-1 in Nck-1 Co-IP and Nck1 levels in IRAK-1 Co-ip showing statistical significance after shear stress. **D)** Schematic representation to the pathway activation. **E)** Immunostained ligated left carotid arteries from iEC-Control, Nck1 KO, iEC-Nck2 KO, and iEC-Nck1/2 DKO mice. p-IRAK-1 (red), predominately within the endothelium (vWF, green) is reduced in Nck1 KO mice and **(F)** graphical quantifications showing the significance. Data are from n=4, represented as mean ± SEM, analyzed by 1-Way ANOVA, and Tukey’s post-test, *p<0.05, **p<0.01, ***p<0.001, ****p<0.0001.

### IRAK-1 contributes to early atherogenic endothelial activation

To determine if IRAK-1 contributes to disturbed flow-induced NF-κB activation, IRAK-1 was depleted in HAECs using siRNA SMARTPool treatment (Figure 9F), and the cells were exposed to OSS for 18h. IRAK-1 knockdown significantly blunts NF-κB activation and proinflammatory gene expression (Figure 10 A/B). Finally, we asked Nck1 and phospho-IRAK-1 in endothelial cells overlying early (stage 1) human atherosclerotic plaques. Both Nck1 and phospho-IRAK-1 showed enhanced staining in endothelial cells associated with early atherosclerotic plaques in humans (Figure 10C). Collectively, our data reveals a novel role for Nck1 in endothelial activation by atheroprone hemodynamics and demonstrates a novel link between Nck1 and IRAK-1 activation in mediating disturbed flow-induced endothelial activation (Figure 10D).

**Figure 10.**
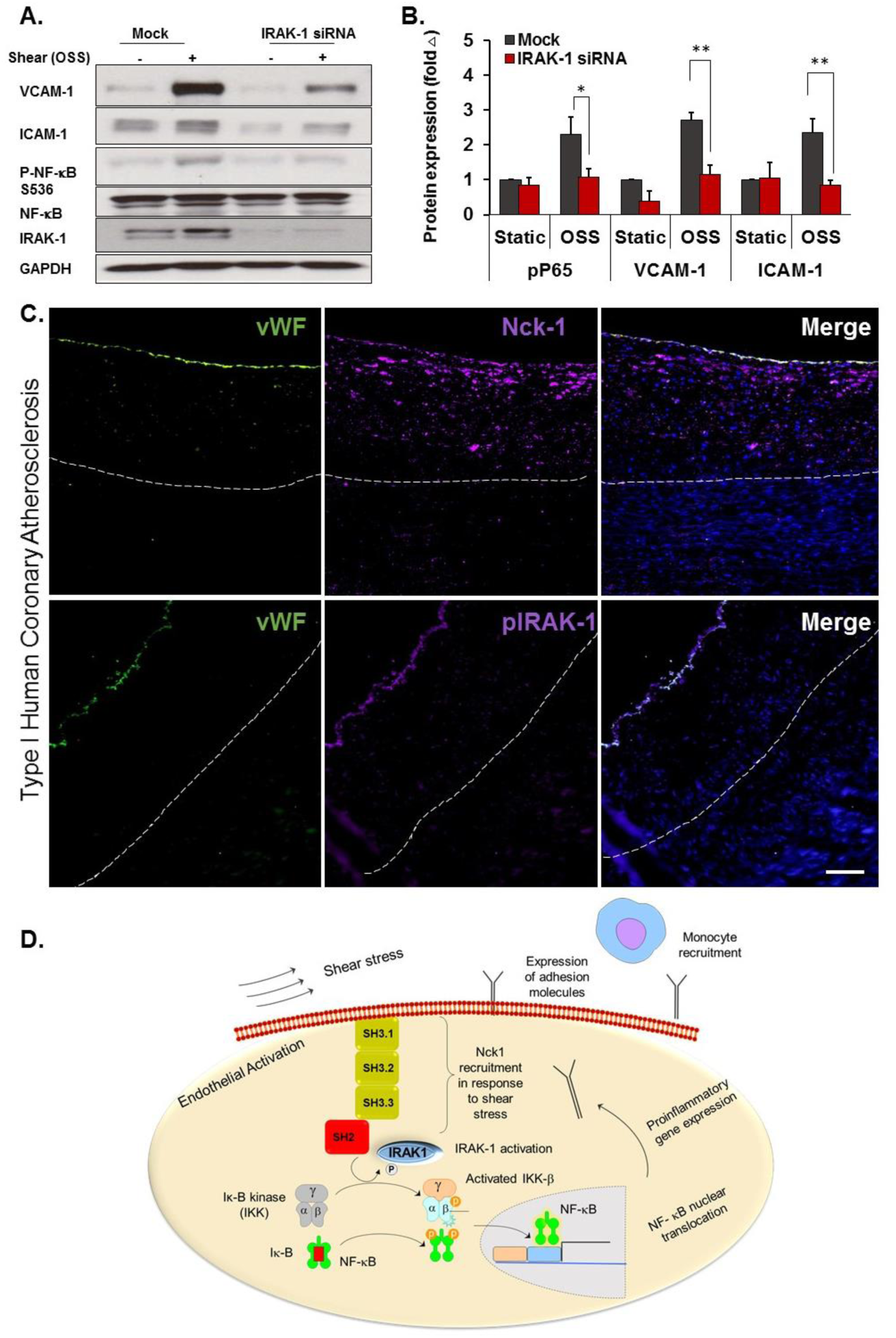
IRAK1 deletion ameliorates shear stress induced activation. **A)** HAECs were treated with or without IRAK-1 siRNA and subjected to OSS (18h). Cell lysates were analyzed for VCAM-1, ICAM-1 and phospho-NF-kB using Western blotting. **B)** Quantification of ICAM-1, VCAM-1 and phospho-NF-κB showing significant reduction in IRAK-1 depleted cells. Densitometric analysis was performed using Image j. Data are mean ± SEM, analyzed by 2-Way ANOVA and Bonferroni’s post-test, *p<0.05, **p<0.01. **C)** Type I Human atherosclerotic lesions were stained for p-IRAK-1 and Nck1, and vWF was used as endothelial marker. Scale bar=100µm. Images analyzed using Nis Elements software, from n=5 postmortem biopsy samples. **D)** Schematic of mechanism of Nck1 mediated IRAK-1/ NF-κB activation in endothelial cells. In response to shear stress exposure, Nck1 is recruited to cell membrane. This facilitates IRAK-1 membrane translocation and activation, which in turns, initiates a signaling cascade leading to p65 activation, nuclear translocation, and proinflammatory gene expression. The subsequent expression of adhesion molecules promotes monocyte recruitment to initiate atherosclerotic plaque formation.

## Discussion

Atherogenic endothelial activation promotes vascular permeability and enhances adhesiveness for circulating leukocytes (28). In early atherosclerosis, atheroprone disturbed flow (low magnitude blood flow with complex features including turbulence, oscillations, separation, and reattachment) promoting endothelial activation (2). Elucidating the pathogenic mechanism of disturbed flow-induced endothelial activation in early lesion formation may lead to interventions that delay or even prevent lesion progression and complications. In this manuscript, we demonstrate a novel, isoform-specific role for Nck1 in atherogenic endothelial activation *in vitro* and *in vivo*. We show that Nck1 is a critical signaling mediator of disturbed flow-induced NF-κB activation and proinflammatory gene expression, and we demonstrate important roles for the Nck1 SH2 domain and first SH3 domain in mediating this effect. We provide the first validation that IRAK-1 serves as a Nck1-specific binding protein, show a vital role for Nck1 in flow-induced IRAK-1 activation, and demonstrate that IRAK-1 is required for chronic NF-κB activation by disturbed flow. Although highly similar to Nck1, endothelial Nck2 expression is dispensable for flow-induced endothelial activation. *In vivo*, only vascular Nck1 deletion reduces endothelial proinflammatory gene expression, monocyte recruitment, and early plaque formation in both the partial carotid ligation model of disturbed flow and in diet-induced spontaneous atherosclerosis, and both Nck1 and phospho-IRAK-1 show enhanced staining in early human atherosclerotic plaques. Taken together, these data clearly identify a novel isoform-specific role for Nck1 in mediating endothelial activation under atheroprone hemodynamics.

The Nck adaptor proteins have been extensively studied in diverse signaling events, most often affecting pathways leading to cellular morphogenesis (10). Previous studies have identified at least 60 Nck1 and Nck2 associated proteins, mostly involved in cytoskeletal organization (22, 32). In models of cell migration and tissue remodeling, Nck1 and Nck2 appear to be largely redundant. Deletion of both Nck1 and Nck2 during development hinders vasculogenesis, whereas individual deletion of either Nck1 or Nck2 did not (21). In the ischemic retinopathy model, deletion of both Nck1 and Nck2 were required to reduce angiogenesis associated with diminished endothelial and pericyte migration (12, 33). However, Guan *et al.* (34) demonstrated that Nck1 and Nck2 can act independently in dermal fibroblasts’ migration and T cell activation (14). We previously demonstrated that depletion of both Nck1 and Nck2 by siRNA blunts oxidative stress induced NF-κB activation in models of ischemia/reperfusion injury (15), but isoform-specific roles were not addressed. Our current data provide the first direct evidence that Nck1 meditates inflammation due to atheroprone flow and demonstrates an unexpected isoform selective role in mediating disturbed flow-induced NF-κB activation and proinflammatory gene expression within the endothelium.

Composed exclusively of SH2/SH3 domains, Nck proteins have no enzymatic activity but regulate multiple signal transduction pathways by controlling protein localization and signaling complex formation (10, 22). While the modular structure of Nck1/2 allow for numerous individual and probably simultaneous protein-protein interactions (11), our data specifically identify a role for the Nck1 SH2 domain and a redundant role for the first Nck1/2 SH3 domain as the critical sites mediating endothelial activation by atheroprone flow. Although the consensus SH2 binding sequence for Nck1 and Nck2 are highly similar (35), the Nck1 and Nck2 SH2 domains are able to bind distinct phosphotyrosine sequences on growth factor receptors (35) and at sites of cell adhesion (35). Consistent with this selectivity, phosphorylated Nephrin selectively recruits Nck1 and not Nck2 through Nck1’s SH2 domain *in vivo* (30). To our knowledge the binding and signaling properties of individual Nck1 SH3 domains have yet to be systematically explored. The critical SH3.1 domain in Nck1 binds an atypical PxxDY motif that undergoes negative regulation by tyrosine phosphorylation, providing a potential negative feedback response that would be expected to limit atherogenic endothelial activation. However, future studies will be required to identify the Nck1 SH3.1 binding partners important for endothelial activation by atheroprone flow.

To explore the non-compensating role of the Nck1 SH2 and SH3.1 domains in disturbed flow-induced endothelial activation, we sought to identify potential binding partners that could mediate endothelial activation by disturbed flow. A previous report utilizing the BioID proximity biotinylation system to identify Nck1 and Nck2 binding proteins found IRAK-1 as a specific interacting partner for Nck1 (24). While this interaction was proposed to be SH2-dependent using computer modeling software, the specific domain mediating this interaction was not determined. Our Co-IPs confirm the direct interactions between IRAK-1 and Nck1, but not Nck2, and demonstrate enhanced interactions following exposure to disturbed flow.

IRAK-1 was first identified through its association with IL-1 receptors at plasma membrane (31) and is a critical mediator of IL-1-induced NF-κB activation (36). Ubiquitously expressed, IRAK-1 has been extensively studied in immune cells (37) and is activated in response to Toll like receptor (TLR) and IL-1 receptor (IL-1R) signaling. Interestingly, IRAK-1 activation is critically dependent on T209 phosphorylation (38), and T209 was highly phosphorylated in HAECs exposed to OSS. Activation of TLRs and IL-1Rs induces binding of MyD88, a signaling scaffold that recruits IRAK-1 to promote its maximum activation and couple it to downstream signaling partners (36). While Nck1 and MyD88 have disparate domain structures, Nck1 may similarly mediate IRAK-1 recruitment to activated cell surface receptors and facilitate its coupling to downstream signaling components to activate NF-κB by disturbed flow. Indeed, phospho-IRAK-1 was reduced when Nck1 was deleted in our *in vitro* and *in vivo* systems, suggesting that in endothelial cells Nck1 is an upstream mediator of IRAK-1 activation. Significantly, phospho-IRAK-1 and Nck1 show enhanced staining within the endothelium in early human atherosclerosis. In line with this finding, a genome wide association study in 34,541 coronary artery disease patients of European biobanks identified Nck1 as a novel coronary artery disease susceptibility loci (18). These data provide the first description of Nck1’s potential role in this regard.

Due to its known role in vasculagenesis and angiogenesis, Nck1/2 signaling was previously viewed as a poor target to treat atherosclerotic cardiovascular disease. A blocking peptide to the Nck1/2 SH3.2 (second SH3) domain reduces angiogenesis *in vivo* (39), blunts vascular permeability in models of I/R injury (15) and atherosclerosis (40), and limits inflammation in atherosclerosis and LPS-induced lung injury^39,40^. However, targeting both Nck1 and Nck2 with this inhibitor is unlikely to be beneficial toatherosclerotic disease, as the inhibitor would limit angiogenesis in ischemic regions affected by the plaque. Our findings suggest the possibility of selectively targeting Nck1 to limit plaque associated inflammation without hindering angiogenesis in ischemic tissue, as Nck2 can compensate for Nck1 in this regard. Since inhibition of either isoform alone is not sufficient to reduce angiogenesis (12, 33), targeting Nck1 may represent a realistic therapeutic to limit endothelial activation without affecting angiogenic tissue remodeling.

In conclusion, our results provide the first data linking endothelial Nck1 signaling to atherogenic endothelial activation and atherosclerotic plaque development. Furthermore, we identified the Nck1 SH2 domain and IRAK-1 as critical mediators of atheroprone flow-induced NF-κB activation and proinflammatory gene expression. Taken together, our findings extend our current understanding of endothelial cell activation in response to atheroprone hemodynamics and identify inhibition of Nck1 a potential future therapeutic in atherosclerotic cardiovascular disease.

## Methods

### Cell culture, Plasmids and RNA interference

Human aortic endothelial cells (HAECs, CELL Applications) were maintained in MCDB131 containing 10% (v/v) fetal bovine serum and supplemented with bovine brain extract (24 µg/ml), 2 mmol/L glutamine, 10 U/ml penicillin, and 100 µg/ml streptomycin. Cells were used between passages 6-10. HAEC transformation was induced by transduction of HAECs with a lentiviral vector expressing SV40-T as previously reported (41). Mouse aortic endothelial cells (MAECs) from Nck1^+/+^ Nck2^fl/fl^ and Nck1^−/−^ Nck2^fl/fl^ (gift of Dr. Tony Pawson, Samuel Lunefeld Research Institute) were isolated from aortic rings as previously described (42) and the cells were kept in MCDB131 containing 10% (v/v) media. To induce Nck2 or double knock outs, the cells were infected with adenovirus expressing either GFP (1×10^8^ PFU/ml) or GFP-2A-iCre (improved mammalian expression, conjugated to GFP, 2×10^8^ PFU/ml; VECTOR BIOLABS) and sorted for GFP positivity. For shRNA delivery, the lentiviral vector used was pLV-(shRNA)-mCherry:T2A:Puro-U6 (Nck1 target seq: GGGTTCTCTGTCAGAGAAA; Nck2 target seq: CTTAAAGCGTCAGGGAAGA; VectorBuillder) with 3^rd^ generation lenti components provided from Addgene; pMD2.G (12259), pRSV-Rev (12253), pMDLg/pRRE (12251), all are deposited from Didier Trono (43). The resulting plasmids were used to package lentiviruses and infect target cells. Transfection with SMARTpool siRNAs targeting Nck1 (L-006354) and Nck2 (L-019547; IDT) was performed using Lipofectamine 3000 (Invitrogen) according to the manufacture’s recommendations. CRISPR/Cas9 deletion of Nck1 and Nck2 utilized double-guide RNA (dgRNA) sequences targeting Nck1 (GTCGTCAATAACCTAAATAC) Nck2 (TGACGCGCGACCCCTTCACC), or Scrambled sgRNA (GCACTACCAGAGCTAACTCA). The lentiviral vectors encoding the full length Nck1 and Nck2 were generated in pLV-EXP-mCherry:T2A:Puro-CMV>Nck vector. The domain swap experiments utilized pLV-EXP-mCherry:T2A:Puro-CMV>Nck constructs containing Nck1 SH2 and Nck2 SH3 domains or Nck2 SH2 and Nck1 SH3 domains. Point mutations in Nck1 SH2 and SH3 domains were conducted by the Redox Molecular Signaling Core (COBRE Center for Cardiovascular Diseases and Sciences, LSUHSC, LA, USA). Nck1 domains point mutations have been verified by cloning PCR and sequencing (Eurofins Genomics) before being used for experiments.

### Induction of acute onset and chronic shear stress in vitro

Shear stress experiments were performed using the parallel plate flow chambers as we previously published (44, 45). To test the early onset of the proinflammatory endothelial activation, cells were subjected to acute shear stress (12 dynes/cm^2^) for up to 45 minutes. Static cell culture was used as a negative control. Oscillatory shear stress (OSS) was induced using a syringe pump (±5 dynes/cm^2^ with a superimposed 1 dynes/cm^2^ for waste exchange) for 18 hours. Chronic laminar shear stress was induced by subjecting the cells to 12 dynes/cm^2^ for 18 hours. After cessation of flow, the cells were rapidly lyzed using 2X Laemmli buffer for Western blot, TRIzol for qRT-PCR, or fixed in 4% (v/v) formaldehyde for immunostaining.

### Western Blot

Proteins in the cell lysates were separated using SDS-PAGE gel and transferred to PVDF membranes (BioRad). After blocking the non-specific bindings with 5% (w/v) non-fat dry milk in 0.1% (v/v) TBS-Tween-20, the membranes were incubated with primary antibodies (See Table I in online Data Supplements for details) at 4°C overnight. After secondary antibody incubation and chemiluminescence, densitometry was analyzed using NIH Image J software.

### Quantitative real time PCR

Total RNAs were extracted using TRIzol® (Thermofisher) from mouse carotid arteries (described below) and HAECs of Nck1 and Nck2 knockdown and knockout cells. Reverse transcription was performed with iScript kit (Bio-rad, 1708890). qRT-PCR was performed using SYBR® Green Master Mix (Bio-rad, 1708882), and gene expression was quantified using the ddCt method. The gene expression was normalized to RPL13a and B2M (Table II lists the primers used in the study, Online Data Supplements).

### Mouse experiments

The mice were housed at LSUHSC-animal care facility under standard 12:12-h light: dark cycle and fed with standard rodent chow and water *ad libitum*. Sample size was determined for each experiment. ApoE^−/−^ mice on the C57BI/6J background were purchased from Jackson Laboratory (Bar Harvor, ME). Mice containing alleles for global Nck1 knockout (Nck1^−/−^) and conditional Nck2 deletion (Nck2^fl/fl^) were a gift from Dr. Tony Pawson, and mice containing the vascular endothelial (VE)-cadherin-driven tamoxifen-inducible Cre (VE-Cadherin CreERT2) were kindly provided by Dr Luisa Iruela-Arispe, (UCLA). Mice were crossed with ApoE^−/−^ to produce inducible, endothelial specific (iEC)-Control mice (ApoE^−/−^, VE-cadherin CreERT2^tg/?^), Nck1 KO mice (ApoE^−/−^, VE-cadherin CreERT2^tg/?^, Nck1^−/−^), iEC-Nck2 KO mice (ApoE^−/−^, VE-cadherin CreERT2^tg/?^, Nck2^fl/fl^), and iEC-Nck1/2 DKO mice (ApoE^−/−^, VE-cadherin CreERT2^tg/?^, Nck2^fl/fl^, Nck1^−/−^). Male mice at 8-10-week old were intraperitoneally injected with Tamoxifen (1mg/kg, Sigma, St Louis, MO) for five consecutive days to induce Cre nuclear translocation and gene excision. After 2 week recovery, the four groups of animals were either subjected to partial carotid ligation (PCL) surgery as we previously reported (25) or fed high fat diet (TD 88137, Harlan-Teklad, Madison, WI) for 12 weeks to induce spontaneous atherosclerotic plaque formation. For PCL, after induction of anesthesia (4% v/v Isoflurane/O_2_), the left external carotid (below the superior thyroid artery), internal carotid and occipital arteries were ligated with 6-0 sutures, whereas the right carotid artery was left unligated and served as an internal control. Disturbed flow within the left common carotid artery below the ligation was confirmed using Doppler Ultrasound (VisualSonics VEVO3100 System) as previously described (26). Mice were euthanized either 48h post-ligation surgery for TRizol flush and intimal/ medial and adventitial mRNA isolation or 7 days post-ligation surgery for immunohistochemistry.

For bone marrow transplant experiments, Nck1 WT or Nck1 KO mice at 8-10 weeks were irradiated with 2 cycles of irradiation (525 rads each cycle) and then received 2×10^6^ bone marrow cells by retro-orbital injection from donor mice (Nck1 WT or Nck1 KO). The animals were maintained on 0.2% (w/v) Neomycin sulfate (Sigma) in their water for 2 weeks and at 4 weeks post-radiation, four groups of mice received HFD for additional 12 weeks.

For atherosclerosis studies, mice were fed high fat diet and their body weights were monitored weekly. After the duration of the study (12 weeks), mice were euthanized by pneumothorax under anesthesia (isoflurane/O_2_ 4% (v/v)) and blood was collected by inferior vena cava puncture into heparinized collecting tubes. Plasma was collected following centrifugation at 500 x g for 5 minutes. Tissues were collected and stored in 10% (v/v) Formalin until analysis.

### Lipid analysis

Plasma total cholesterol, high density lipoprotein and triglycerides were analyzed using commercially provided kits as previously described (46).

### Assessment of atherosclerotic lesions

The extent of atherosclerosis was assessed in the whole aorta by an *en face* method (41, 47). Lesion areas were analyzed using NIS Elements and quantified as % of the total surface area. A second assessment to atherosclerosis was conducted in cross-sectional aortic roots, right carotid sinus, left carotid sinus, and brachiocephalic arteries as described (41).

### Immunostaining

For immunofluorescent immunocytochemistry (IF-C), cells were fixed in 4% (v/v) formaldehyde/PBS for 10 minutes and then permeabilized with 0.1% (v/v) Triton X-100 for 5 minutes. After blocking (10% (v/v) Horse serum/1% (v/v) BSA / PBS for 1h), the cells were stained with corresponding primary antibody (Table I, IF-C). For nuclear p65 quantification, at least 100 cells were counted from at least 3 random fields per experimental condition from four independent experiments. For immunohistochemistry (IHC-IF), tissue was fixed with 10% (v/v) formalin, paraffin embedded and sectioned into 5-µm sections as we previously published (41). Heat mediated antigen retrieval pretreatment using 10mM Sodium Citrate buffer (Vectors Biolabs) was used. All primary antibodies (Table I, IHC-IF) were incubated at 4 °C overnight. Images were captured with a Nikon microscope and analyzed using Nis Elements software. The investigators were blinded to the animal groups during the process of data collection and analysis.

### Plasma Cytokine analysis

Freshly isolated plasma was rapidly collected as previously published (46) and cytokine levels were assessed using LEGENDplex™ (BioLegend, cat. No. 740446) according the manufacture’s recommendations. Data analysis was performed using LEGENDplex v8.0 software.

All reagents were provided from Gibco, USA, unless otherwise stated. All lentiviral vectors were designed and obtained using the VectorBuilder website, and site-directed mutagenesis performed by the COBRE Redox Molecular Signaling Core.

### Statistics

Data are analyzed as mean ± standard error of the mean (SEM) using GraphPad prism software (Version 7, GraphPad, San Diego, CA). Data was first tested for the Normality using Kolmogorvo-Smirnov test and then for multiple comparisons, 1-Way ANOVA followed by Tukey’s post-test or 2-Way ANOVA and Bonferroni’s post-test was performed for the normally distributed data. Statistical significance was achieved when p<0.05.

### Study approval

All animal work was performed according to the National Research Council’s Guide for the Care and Use of Laboratory Animals and were approved by LSU Health-Shreveport Institutional Animal Care and Use Committee. Studies involving the use of human atherosclerosis autopsy specimens were approved by the institutional review board of the LSU Health-Shreveport.

## Author contributions

MA performed experiments, data collection, interpretation of data and analysis, manuscript co-writing; DDW, CA, JGT, atherosclerosis experiments’ tissue collection and analysis; AWO, directed the project, designed the study, interpreted results, and co-wrote the manuscript.

## Acknowledgements

This work was supported by a Malcolm Feist Postdoctoral Fellowship and an American Heart Association Postdoctoral Fellowship [AHA grant number 20POST35120288] to MA and by National Institutes of Health grants [HL098435, HL133497, HL141155, and GM121307] to AWO. The authors thank the late Dr. Tony Pawson (Lunenfeld-Tanenbaum Research Institute, Univ. of Toronto) for providing the Nck1^−/−^ and Nck2^flox/flox^ mice), and Dr. Luisa Iruela-Arispe (UCLA, CA) for providing VeCadherin-CreERT2 mice. The authors acknowledge the COBRE Center for Redox Biology and Cardiovascular Disease and the Redox Molecular Signaling Core [P20GM121307] for site-directed mutagenesis of Nck1 constructs.

