## Supplemental materials for "Selective role of Nck1 in atherogenic inflammation and plaque formation"

### SUPPLEMENTAL DATA:

#### Supplemental Tables:

##### *Supplemental Table I*

| Target | Source | Catalog Number | Application | Working Concentration |
|---|---|---|---|---|
| **Phospho-NF-κB (Ser536, p65 subunit)** | Cell Signaling Technology | 3033 | WB | 1:1000 |
| **NF-κB (p65 subunit)** | Cell Signaling Technology | 4764 | WB | 1:1000 |
| **NF-κB (p65 subunit)** | Cell Signaling Technology | 6956 | IF-C | 1:800 |
| **Nck1** | Cell Signaling Technology | 2319 | WB | 1:1000 |
| **Nckb** | Abcam | Ab109239 | WB | 0.45 μg/ml |
| **Nck1/2** | Millipore | 06-288 | WB | 0.1 μg/ml |
| **VCAM-1** | Abcam | Ab134047 | WB | 0.44 μg/ml |
| **GAPDH** | Cell Signaling Technology | 2118 | WB | 1: 5000 |
| **von Willibrand Factor** | Abcam | Ab11713 | IHC-IF | 0.3 μg/ml |
| **ICAM-1 (YN1/1.7.4)** | Abcam | Ab119871 | IHC-IF | 0.25 μg/ml |
| **ICAM-1** | Santa Cruz | sc-1511 | WB | 0.5 μg/ml |
| **Mac2** | Accurate Chemical | CL8942AP | IHC-IF | 0.1 μg/mL |
| **Smooth Muscle Actin** | Sigma Aldrich | C6198 | IHC-IF | 0.25 μg/ml |
| **Phospho-eNOS (Ser1177)** | Cell Signaling Technology | 9571 | WB | 1:500 |
| **Anti-eNOS/NOS Type III** | BD Transduction Lab | 610297 | WB | 0.25μg/ml |
| **Phospho-Akt (Ser473)** | Cell Signaling Technology | 4060 | WB | 1:1000 |
| **AKT1/2 (N-19)** | Santa Cruz | sc-1619 | WB | 0.2 μg/ml |
| **ERK1** | Santa Cruz | Sc-94 | WB | 0.1 μg/mL |
| **Phospho-p44/42 MAPK (ERK1/2)** | Cell Signaling Technology | 4377 | WB | 1:1000 |

4 ***Supplemental Table II***

| Gene | Species | Forward | Reverse |
|---|---|---|---|
| **β2-microglobulin** | Mouse | TTCTGGTGCTTGTCTCACTGA | CAGTATGTTCGGCTTCCCATTC |
| **Rpl13a** | Mouse | GGGCAGGTTCTGGTATTGGAT | GGCTCGGAAATGGTAGGGG |
| **β2-microglobulin** | Human | AGCATTCGGGCCGAGATGTCT | CTGCTGGATGACGTGAGTAAA CCT |
| **Rpl13a** | Human | GCCATCGTGGCTAAACAGGTA | GTTGGTGTTCATCCGCTTGC |
| **VCAM-1** | Mouse | TCAAAGAAAGGGAGACTG | GCTGGAGAACTTCATTATC |
| **VCAM-1** | Human | ATGAGGGGACCACATCTACG | CACCTGGATTCCTTTTTCCA |
| **ICAM-1** | Human | TGTCCCCCTCAAAAGTCATC | TAGGCAACGGGGTCTCTATG |
| **ICAM-1** | Mouse | CTGGCTGTCACAGAACAGGA | AAAGTAGGTGGGGAGGTGCT |
| **KLF2** | Mouse | AGAATGCACCTGAGCCTGCTAG | AATTTCCCCGAAAGCCTGC |
| **SMA** | Mouse | GGACGTACAACTGGTATTGTGC | CGGCAGTAGTCACGAAGGAAT |
| **Nck2** | Mouse | GTCATAGCCAAGTGGGACTACA | GCACGTAGCCTGTCCTGTT |
| **CD31** | Mouse | GGAGTCAGAACCCATCAGGA | CAGCTGGTCCCCTTCTATGA |

***Supplemental Table III: Genotyping Primers:***

| Gene | Sequence |
|---|---|
| **Nck1** | GCATGTAGACAATTACACTTC AGC ACC |
| | ATTCATGGAATTTCGAACTCGCCACC |
| | CTGATTGAAGCAGAAGCCTGCGATG |
| | TATTGGCTTCATCCACCACATACAGG |
| **APOE** | GCCGCCCCGACTGCATCT |
| | TGTGACTTGGGAGCTCTGCAGC |
| | GCCTAGCCGAGGGAGAGCCG |
| **VeCad Cre** | ACT GGG ATC TTC GAA CTC TTT GGA C |
| | GAT GTT GGG GCA CTG CTC ATT CAC C |
| | CCA TCT GCC ACC AGC CAG |
| | TCG CCA TCT TCC AGC AGG |
| **Nck2** | GGA TAC CAC CAT TGG CAT TAG TAG |
| | GTG CTC ATT TGA CAA GTG ACA C |

#### Supplemental Figures:

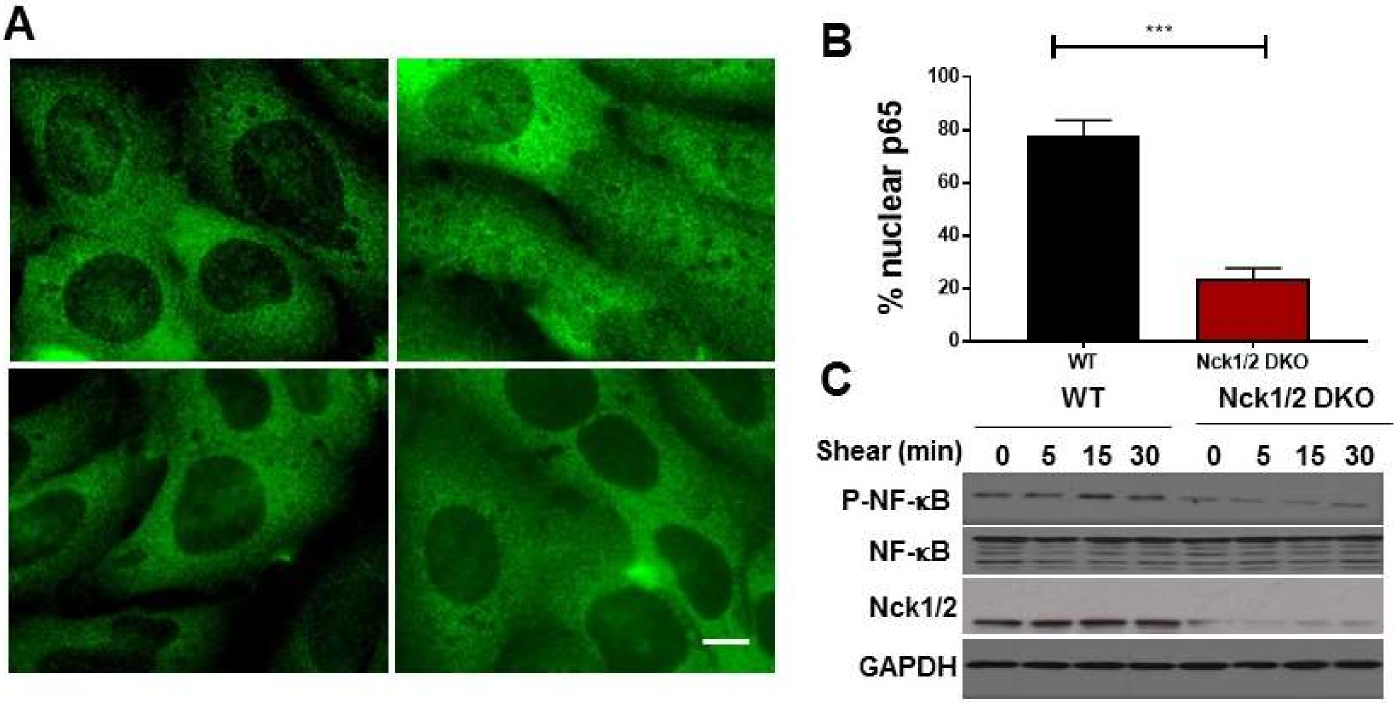

**Supplemental Figure 1. Mouse aortic endothelial cells (MAECs) from iEC-Nck1/2 DKO were isolated and subjected to shear stress. A-C)** NF-κB activation was assessed by analysis of p65 nuclear translocation and **(C)** p65 Ser536 phosphorylation, showing significant reduction in Nck1/2 DKO MAECs compared to Wild-type cells (WT). Data are mean ± SEM, analyzed by unpaired student *t* test, n=4, ***p<0.001. Scale bar=50μm.

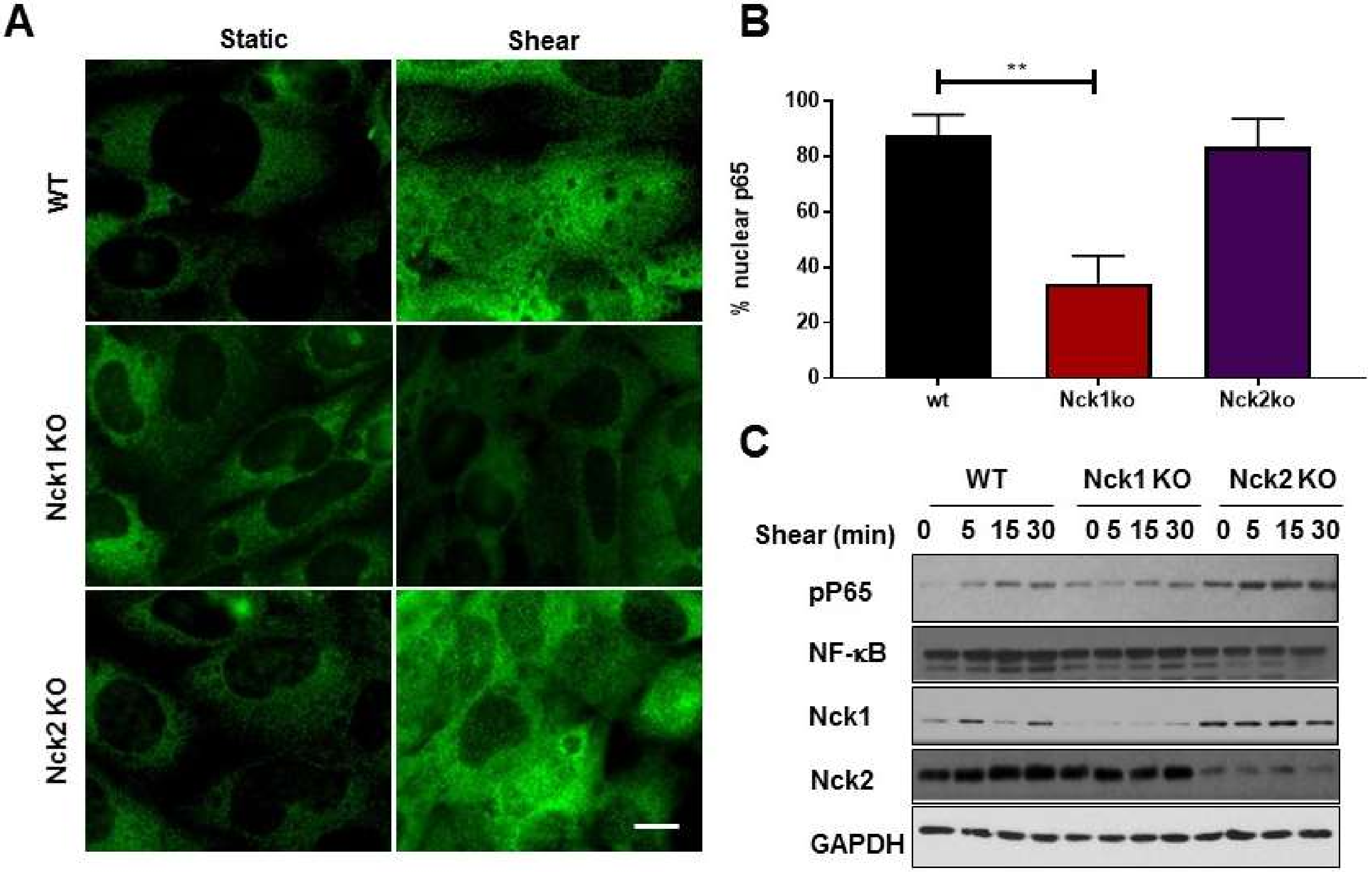

**Supplemental Figure 2. Mouse aortic endothelial cells (MAECs) from Nck1 KO and iEC-Nck2 KO were isolated and subjected to shear stress. A-B)** NF-κB activation was assessed by analysis of p65 nuclear translocation and **(C)** p65 Ser536 phosphorylation, showing significant reduction in Nck1 KO MAECs compared to Wild-type cells (WT). Data are mean ± SEM, analyzed by 1-Way ANOVA, and Tukey's post-test, n=4, **p<0.01. Scale bar=50μm.

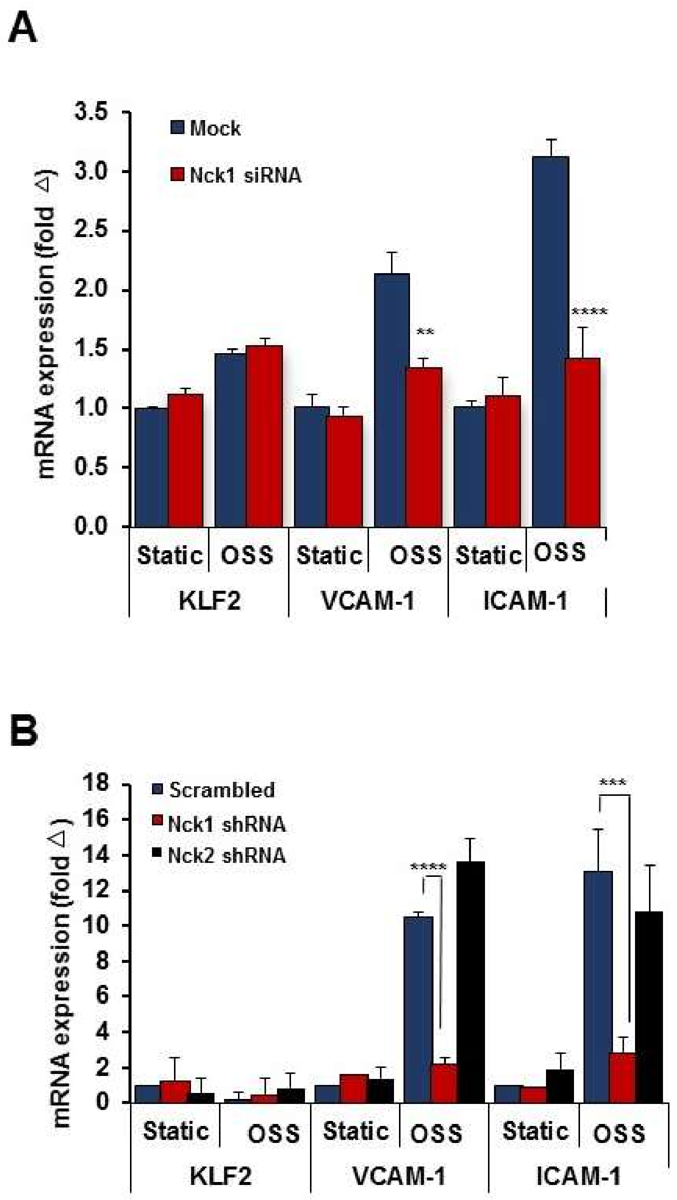

**Supplemental Figure 3. A)** mRNA expression for KLF2, VCAM-1 and ICAM-1 assessed in human aortic endothelial cells (HAECs) knocked down for Nck1 and then subjected to oscillatory shear stress (+/- 5dynes/cm$^2$ with 1 dynes/cm$^2$ for forward flow) for 18h. **B)** mRNA expression of KLF2, VCAM-1 and ICAM-1 in shRNA treated cells and subjected to OSS. Data are from n=4 independent experiments, mean ± SEM, analyzed by 2-Way ANOVA followed by Bonferroni's post-test, **p<0.01, ***p<0.001, ****p<0.0001.

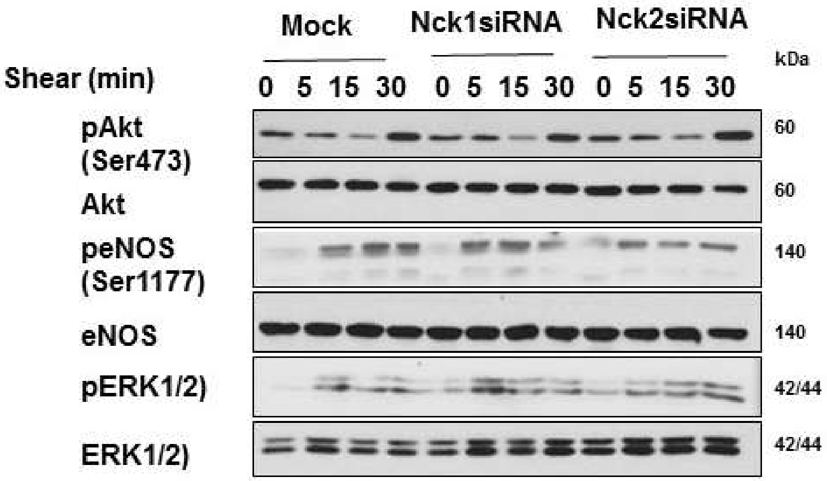

**Supplemental Figure 4.** Endothelial cells subjected to acute shear stress for the indicated times, and activation of Akt, eNOS, and ERK1/2 assessed by Western blotting. Representative blots are from n=4.

.

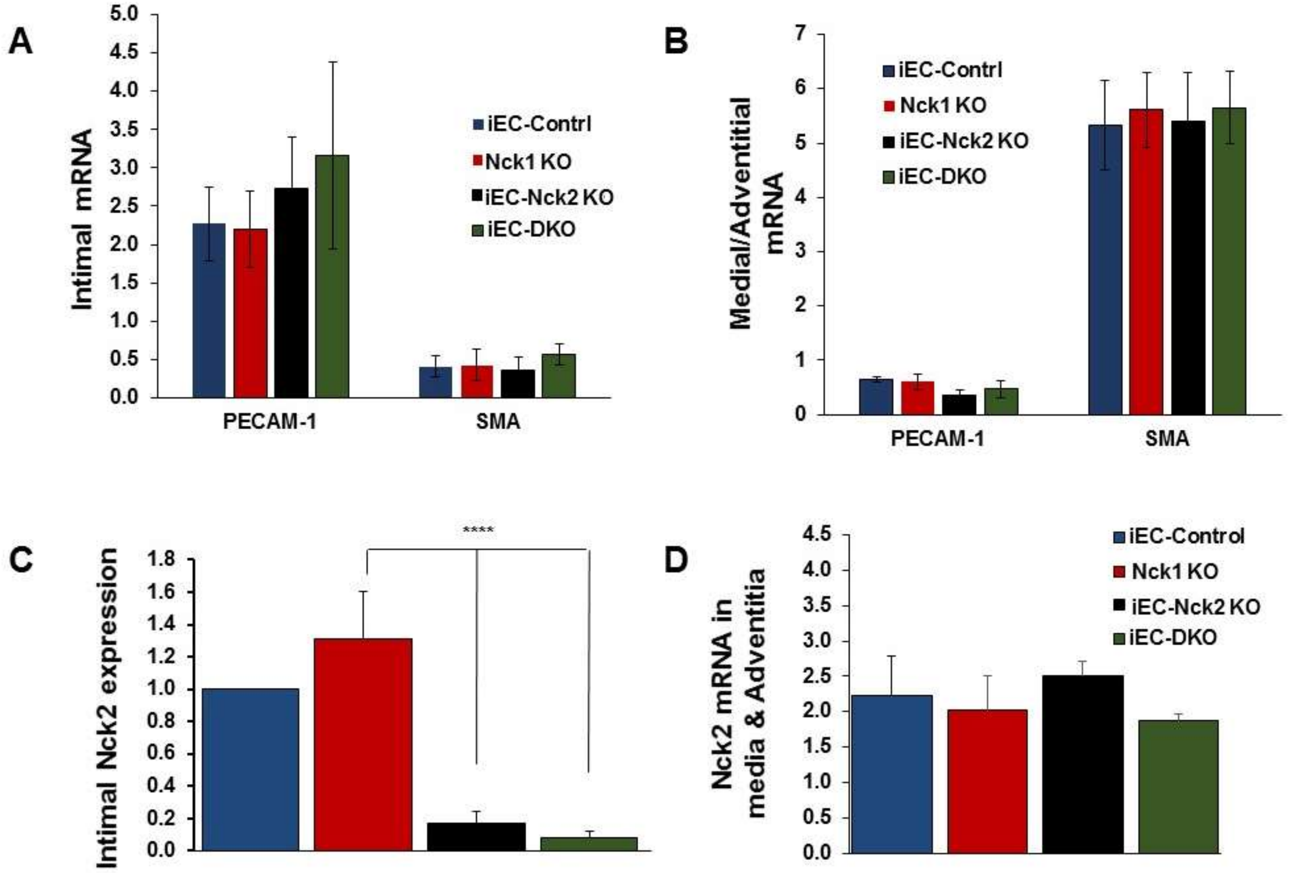

**Supplemental Figure 5. A/B)** Intimal mRNA and Medial/Adventitial mRNA were obtained 2 days post-ligation surgery from right and ligated left carotid arteries. mRNA levels were assessed for the endothelial marker gene platelet endothelial cell adhesion molecule-1 (PECAM-1) and the smooth muscle marker gene α-smooth muscle actin (SMA). **C/D)** Nck2 mRNA expression after tamoxifen injection shows selective Nck2 depletion from the intima and not the media/adventitia. Graphs are mean ± SEM, n=7-10/group. Data analyzed by 1-Way ANOVA and Tukey's post-test, ****$p<0.0001$.

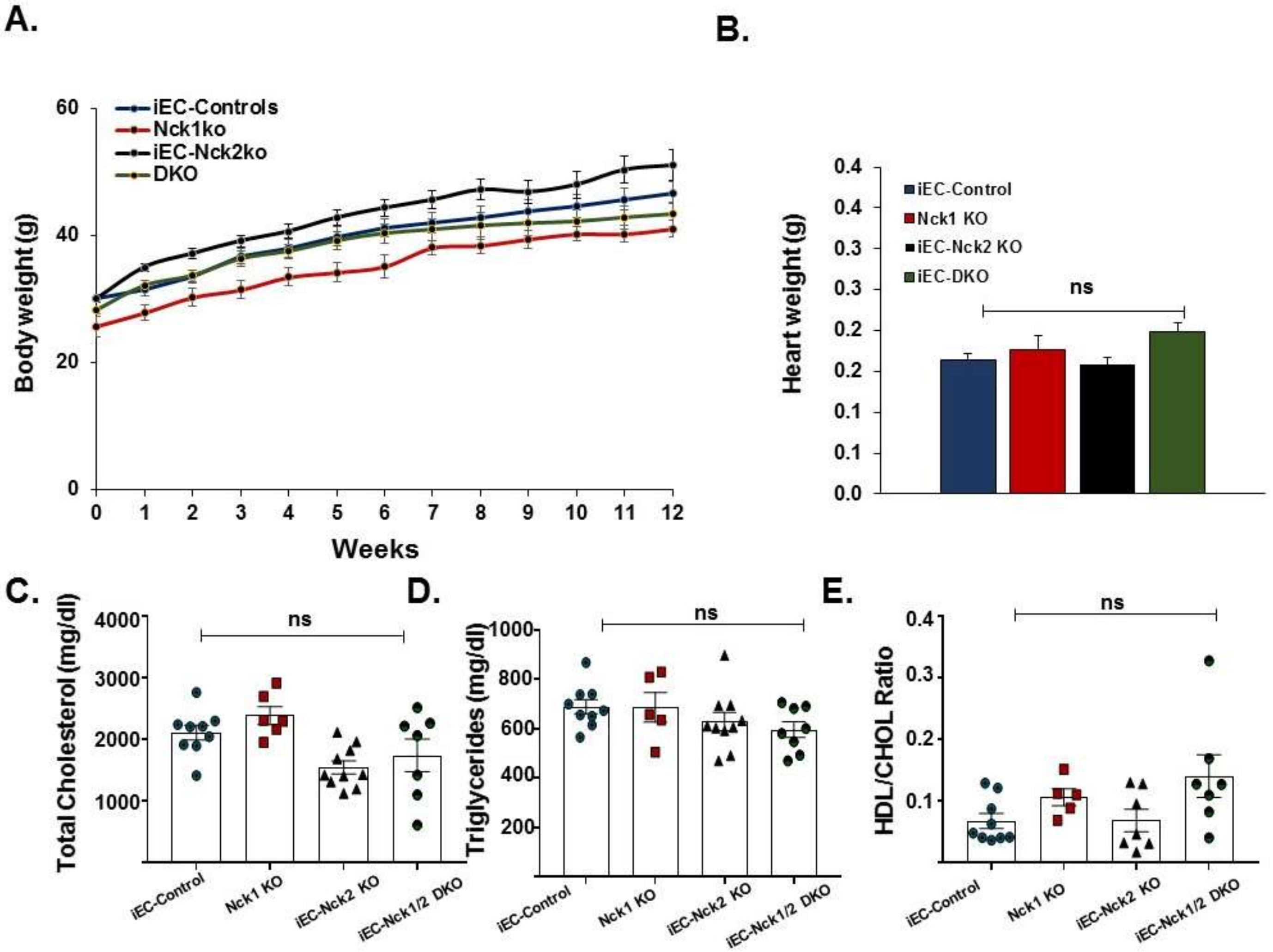

**Supplemental Figure 6. A)** Body weight changes in response to HFD feeding in iEC-Controls, Nck1 KO, iEC-Nck2 KO, iEC-DKO mice. Body weight in grams were recorded weekly for 12 weeks. **B)** Heart weight among experimental groups after 12 weeks of HFD feeding. **C)** Total plasma cholesterol (mg/dl) **(D)** Triglycerides and **(E)** HDL/total Cholesterol ratio were measured. Data are mean ± SEM, n=5-10/group. Data analyzed by 1-Way ANOVA and Tukey's post-test, ns: not-significant.

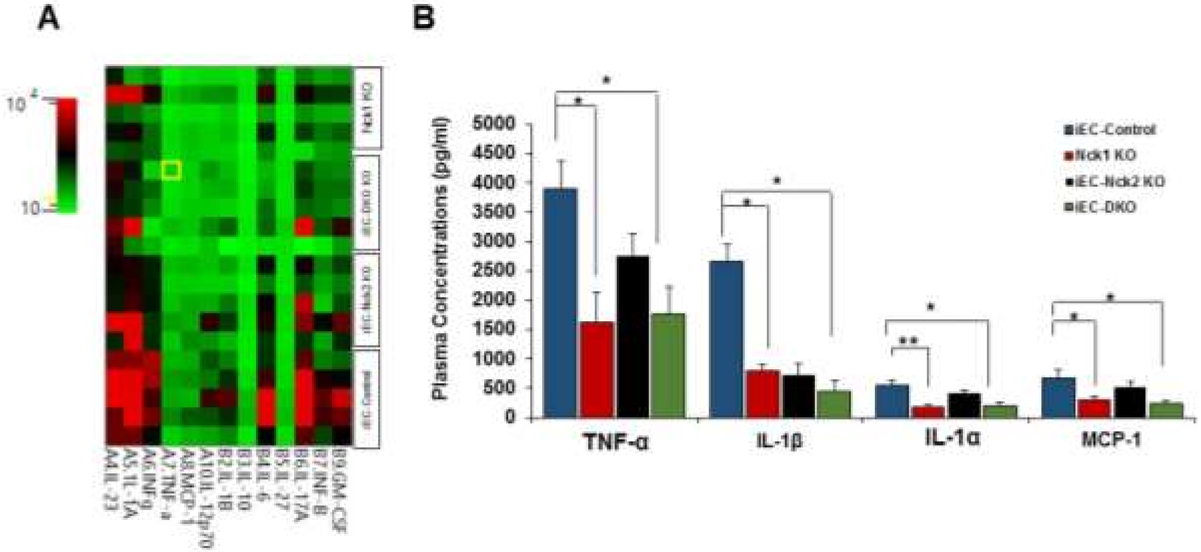

**Supplemental Figure 7. Nck1 KO mice show attenuated plasma pro-inflammatory cytokine/chemokine profiles. A)** Heat map and **(B)** a graphical representation of plasma cytokine concentrations (pg/ml), analyzed using LEGENDplex$^{TM}$. Data are mean ± SEM, n=6-10/group. Data analyzed by 2-Way ANOVA and Tukey's post-test, *p<0.05, **p<0.01.

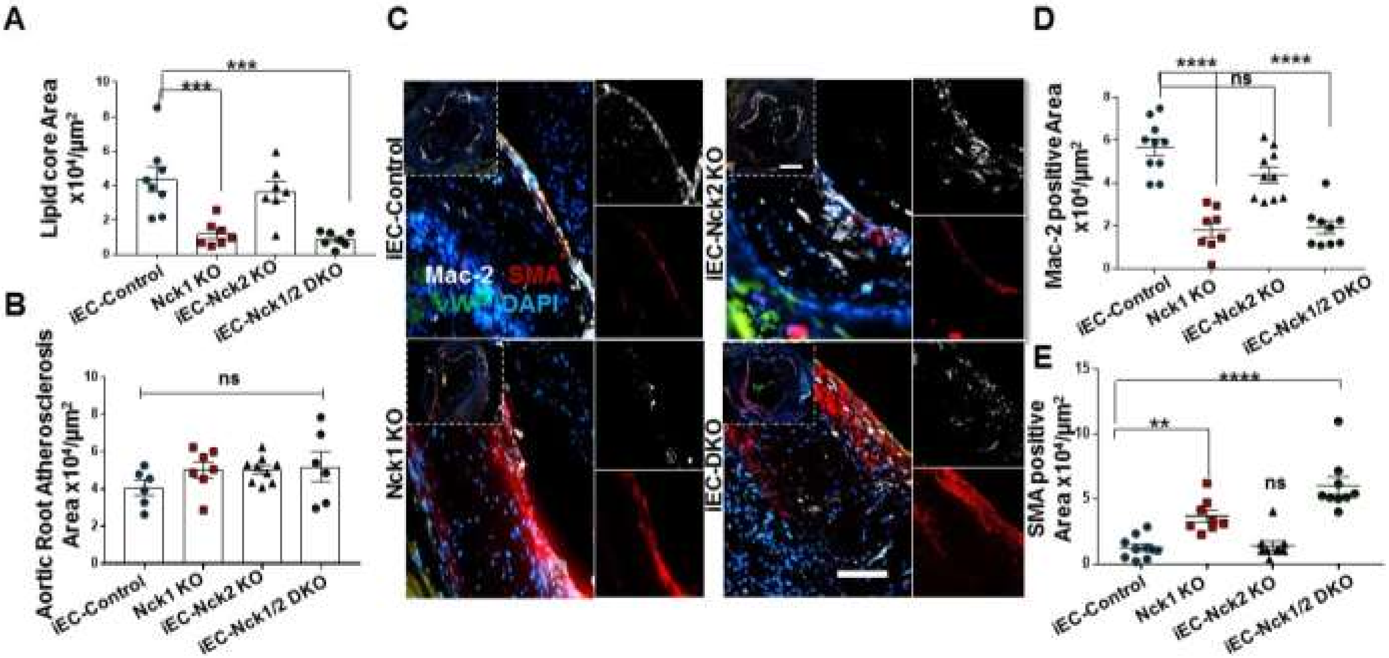

**Supplemental Figure 8. A)** Lipid core area quantification in carotid atherosclerosis. **B)** Aortic Root atherosclerosis and **(C-E)** analysis of aortic root plaque cellular content following staining for macrophages (Mac-2; white), smooth muscle cells (α-smooth muscle actin; SMA, red) and endothelium (vWF; green). Analysis was performed using NIS Elements software and data are mean ± SEM, n=6-10/group. Data analyzed by 1-Way ANOVA and Tukey's post-test, **p<0.01, ***p<0.001, ****p<0.0001. Scale bars=50-200μm, ns=not significant.

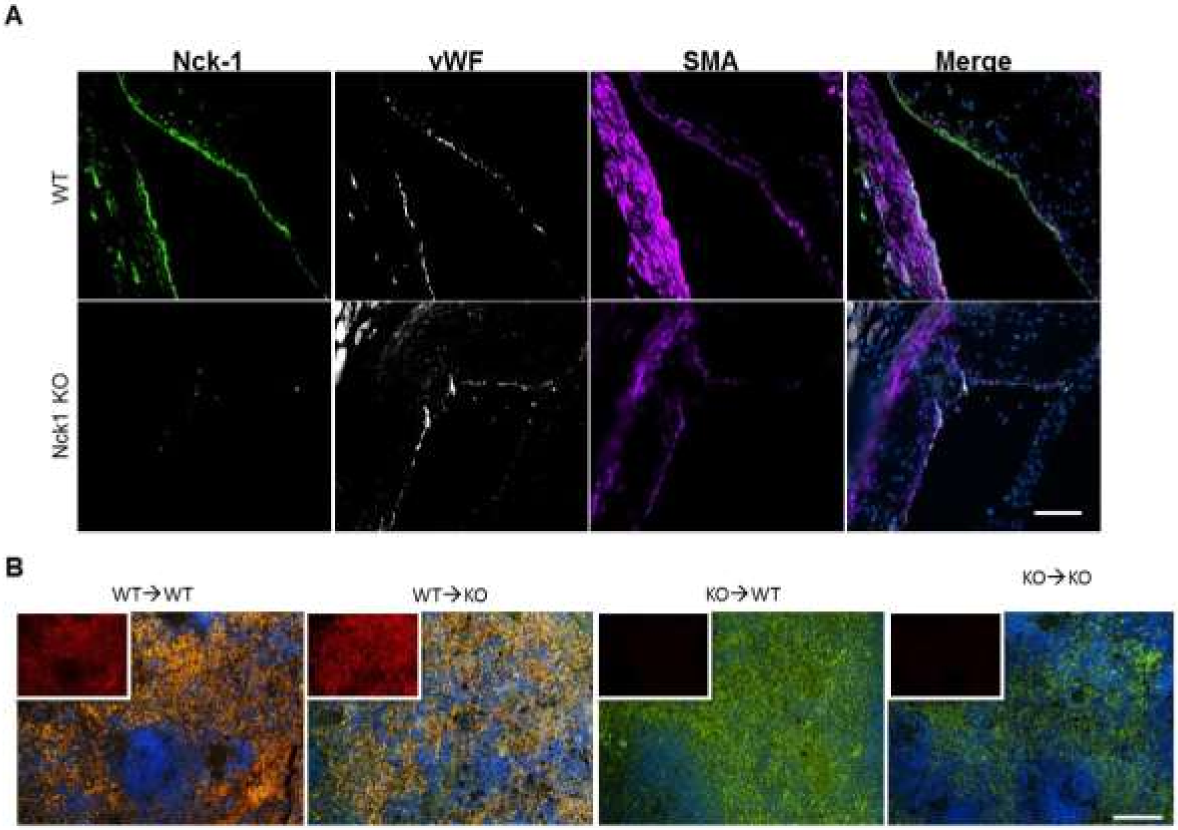

**Supplemental Figure 9. A)** Stained sections of mouse aortic atherosclerosis from wild type (WT) or Nck1 KO mice stained for Nck1, vWF, SMA. **B)** Splenic sections from bone marrow chimeric mice stained for Nck1 (red) and mac-2 (green). From n=5/ group. Scale bars=100μm.

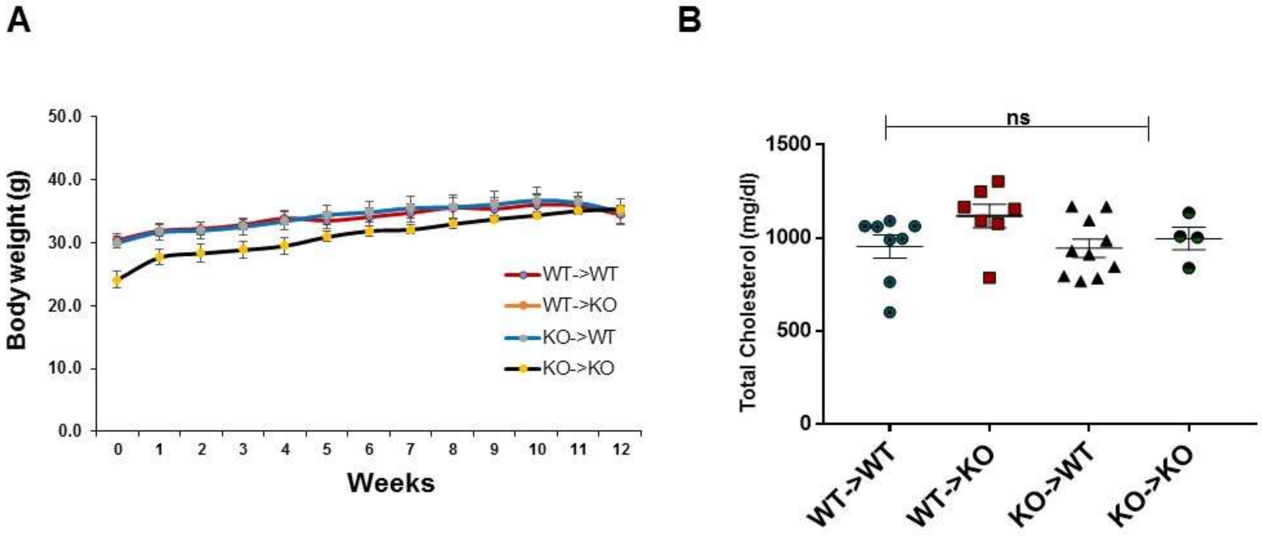

**Supplemental Figure 10.** Bone marrow chimera were produced as described in methods. **A)** Average body weight measurement assessed weekly for 12 weeks during the period of HFD feeding. **B)** Plasma cholesterol levels were assessed using an enzymatic assay. Data are mean ± SEM, 2-Way ANOVA and Bonferroni's post-test was used, ns=not significant.

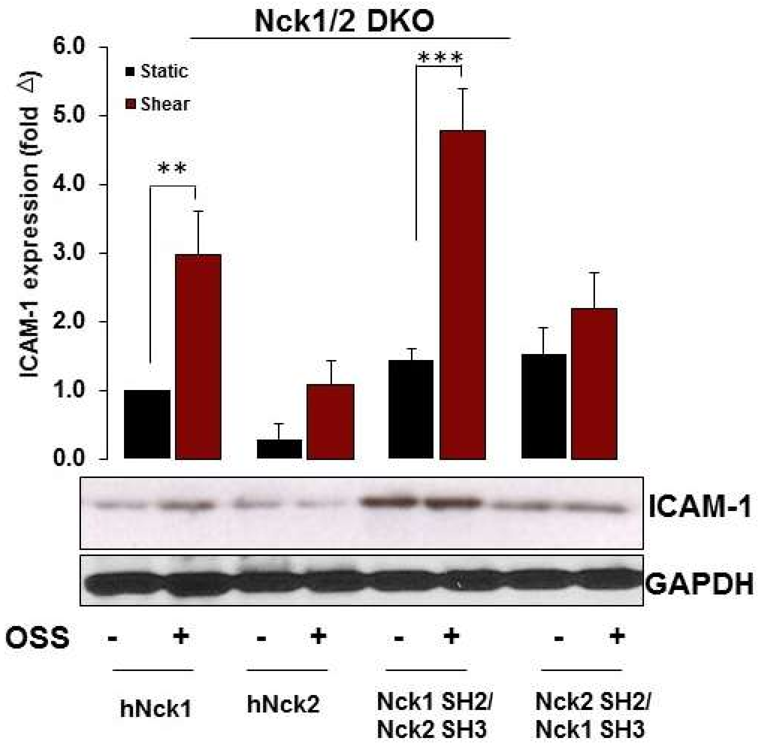

**Supplemental Figure 11.** ICAM-1 protein expression in Nck1/2 DKO HAECs after rescuing the expression of Nck1, Nck2, the Nck1 SH2/Nck2 SH3 chimera, or the Nck2 SH2/Nck1 SH3 chimera. Data are mean ± SEM, n=4 independent experiment. Data are analyzed by 2-Way ANOVA and Bonferroni's post-test, **p<0.01, ***p<0.001.

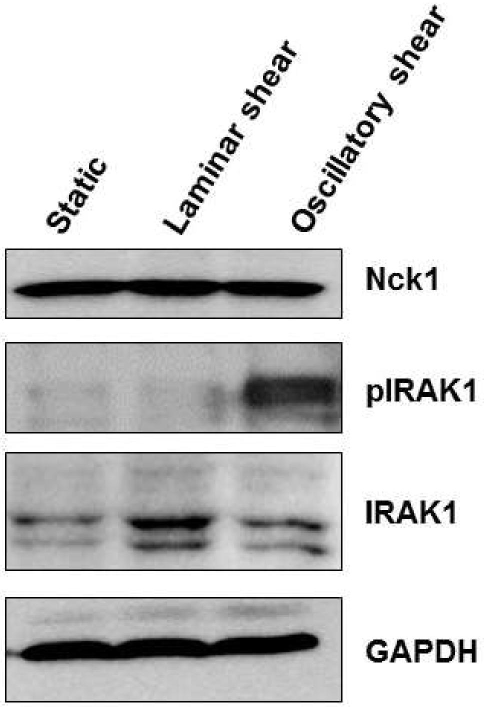

**Supplemental Figure 12.** Human Aortic endothelial cells (HAECs) were subjected to laminar shear stress (10 dynes/cm2, 18h) or oscillatory shear stress (+/- 5dynes/$cm^2$ with 1 dyne/$cm^2$ forward flow, 18h). Cell lysates were assessed using Western blot for Nck1, phospho. IRAK-1, IRAK-1 levels. GAPDH was used a loading control. Representative blots from n=4.

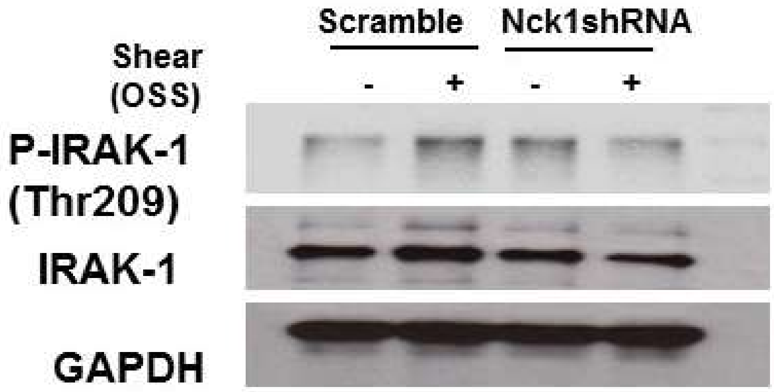

**Supplemental Figure 13.** Lentiviral shRNA transduced **h**uman aortic endothelial cells (HAECs) were subjected to shear stress (+/- 5dynes/cm$^2$ with 1 dyne/cm$^2$ forward flow, 18h). Cell lysates were assessed using Western blot for pIRAK-1, IRAK-1 levels. GAPDH was used a loading control. Representative blots from n=4.

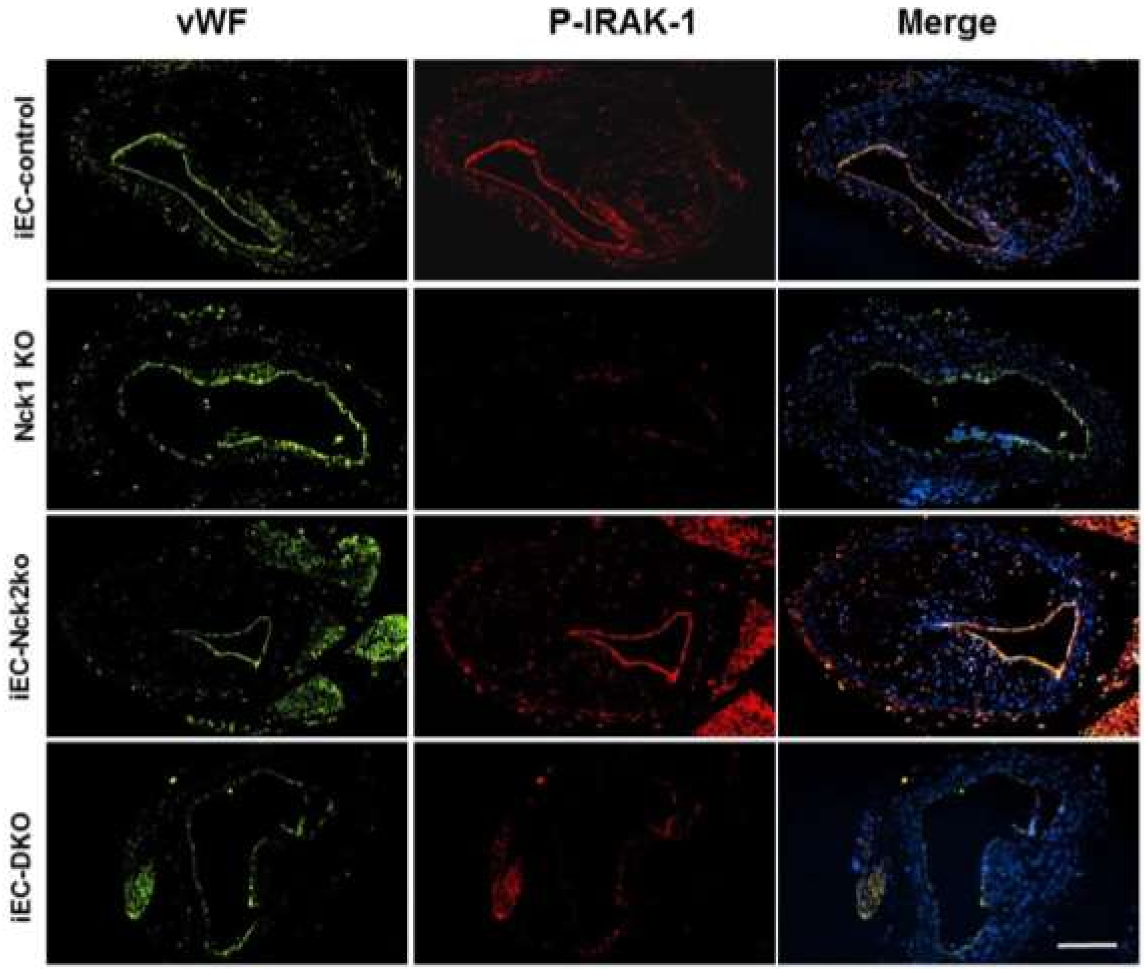

**Supplemental Figure 14.** Immunostained images from brachiocephalic arteries (BCA) after HFD feeding in iEC-Controls, Nck1 KO, iEC-Nck2 KO, iEC-DKO mice. Representative staining for endothelium (vWF; green), p-IRAK-1 (red). Analysis was performed using NIS Elements software and images are from n=4/group. Scale bar=100μm.
